# Mechanistic basis of gene-specific transcription regulation by the Integrator complex

**DOI:** 10.1101/2024.01.24.576984

**Authors:** Kevin Sabath, Amena Nabih, Christian Arnold, Rim Moussa, David Domjan, Judith B. Zaugg, Stefanie Jonas

## Abstract

The Integrator complex (INT) regulates gene expression via premature transcription termination of RNA polymerase II (RNAP2) at promoter-proximal pausing sites. This attenuation of transcription is required for cellular response to external stimuli, cell differentiation and neurodevelopment. How gene-specific regulation is achieved by INT in an inducible manner remains unclear. Here, we identify two sites on INT subunits 13/14 that serve as direct binding hubs for diverse sets of sequence-specific transcription factors (TFs) and other transcription effector complexes. The TFs co-localize with INT genome-wide, increase INT abundance on target genes and co-regulate inducible transcriptional programs. Consistently, disruption of INT-TF contacts impairs sensory cilia formation in response to glucose starvation. Structural analysis places INT’s TF binding hubs upstream of the transcription bubble when attached to paused RNAP2, consistent with simultaneous TF-promoter association. Our data establish TF-mediated recruitment of INT to promoters as a widespread mechanism for targeted and inducible transcription attenuation.

## INTRODUCTION

Accurate gene regulation is essential for cell differentiation during development and spatiotemporal response to external cues. Therefore, as the first step in gene expression, the transcription of mRNAs and non-coding (nc)RNAs by RNA polymerase II (RNAP2) is tightly controlled.^1^ In metazoans, the central decision – for or against production of transcripts – occurs not only at initiation but also during pausing close to transcription start sites (TSS).^2,3^ At this stage, pausing factors (NELF and DSIF) arrest RNAP2 in its catalytic cycle.^4,5^ From this block, RNAP2 can either be released into processive elongation or discharged via premature termination.^6–8^ Elongation requires P-TEFb, which phosphorylates NELF, DSIF, and the C-terminal domain (CTD) of RNAP2.^9–11^ These phosphorylations lead to NELF dissociation, transformation of DSIF into an elongation factor, and recruitment of elongation complexes to RNAP2.^12^ Alternatively, paused RNAP2 (paused-elongation complex, PEC) can also be bound by the Integrator complex (INT),^13–16^ which cleaves the nascent transcript, and keeps NELF, DSIF and RNAP2’s CTD in a dephosphorylated state, thereby counteracting elongation and instead eliciting premature transcription termination.^17^ Thus, INT acts as a repressor of full length mRNA transcription in a process that has been termed attenuation.^18,19^ Furthermore, INT restricts elongation-incompetent RNAP2 complexes from proceeding, which also reduces antisense transcription and production of PROMPTs.^20–22^ Finally, the RNAP2 cleavage and termination functions of INT are also exploited by many essential ncRNAs (e.g. uridine-rich small nuclear (Usn)RNAs, telomerase RNA, piwi RNAs) for the initial 3’ processing of their primary transcripts.^23–27^ INT is a 1.5 MDa complex that consists of 15 subunits (INTS1-15) organized in modules,^28^ and also incorporates two subunits of the general protein phosphatase 2A (PP2Ac, PR65).^29,30^ Structures of INT containing a large fraction of its subunits show that the two catalytic modules – INT RNA cleavage module (ICM=INTS4-9-11) and phosphatase PP2A – sit laterally on two sides of a central scaffold axis.^30^ To specifically recognize paused RNAP2, INT makes contacts with all three components of PEC (RNAP2, DSIF, NELF).^13,16^ As a result of PEC binding, the INTS11 endonuclease becomes activated for cleavage. PP2A on the other hand is thought to prevent or reverse P-TEFb mediated phosphorylations, which are required to initiate elongation.^29–31^

INT-mediated transcription regulation targets many inducible genes with weak promoters.^29,32–34^ Accordingly, INT is not only required for adaptation to diverse stressors such as heat-shock, heavy metal ions, serum starvation and growth factor addition, but also for developmentally important programs such as hematopoiesis, adipogenesis, ciliogenesis, and neuron migration in mouse embryos.^19,32,35–39^ Consistent with a crucial function during cell differentiation and stimulus response, INTS deletions are embryonically lethal and INTS mutations cause developmental disorders, many of which manifest with neurological disease components.^37,40–43^

Despite the clear observation that transcription attenuation via INT is required for diverse biological response pathways, it is unclear how the pivotal decision between productive elongation and abortive early termination is made in the correct spatiotemporal and gene-specific manner. Here, we show that several sequence-specific transcription factors (TFs), and transcription effector complexes (chromatin remodeler INO80; little elongation complex) bind INT directly on two conserved surfaces of the INTS10-13-14 module. The TFs co-regulate gene sets together with INT, co-localize with INT on chromatin and increase INT abundance at promoter-proximal pausing sites of target genes. The TF-interactome of INT is modulated by stress conditions, and TF-INT binding is necessary for primary cilia formation as a cellular response to glucose starvation. Finally, we localize the previously missing INTS10-13-14 module within the INT structure, where it is placed with its TF attachment sites fully accessible and upstream of the transcription bubble, oriented towards the promoter when bound to paused RNAP2. Thus, our data show that INT is recruited to specific genomic loci by TFs in a stimulus responsive manner, where it can efficiently regulate paused RNAP2 close to TSSs.

## RESULTS

### The INTS10-13-14 module binds several sequence-specific transcription factors, the little elongation complex (LEC) and chromatin remodeler INO80

Data from previous studies^35,37^ suggest gene-specific misregulation of inducible transcriptional programs upon depletion or mutation of INTS13, which is part of an INT module consisting of INTS10-13-14.^44^ Therefore, we set out to define the interactome of this module via tandem affinity purification followed by mass spectrometry (TAP-MS) (Figure 1). We used HEK293T-Flp-In cell lines with stably incorporated INTS13 or INTS10 tagged with tandem StrepII-CBP (2SC) (Supplemental Figure 1A, B). To reduce co-purification of indirect binders via other INTS, we deleted the C-terminal 58 amino acids (aa) of INTS13 (2SC-INTS13ΔC), which are required for incorporation of the module into INT by contacts to the ICM.^37,44^ After label-free quantification (LFQ) of MS data, 74 and 76 proteins were significantly enriched in INTS13ΔC and INTS10 TAPs, respectively, compared to TAPs from control cells expressing 2SC-GFP (Figure 1A, B, and Supplemental Figure 1C, D). As expected INTS13ΔC co-purified the other module components INTS10-14, while the INTS10-TAP enriched the entire INT (INTS1-15, PR65, PP2Ac). Notably, INTS15 (previously C7ORF26), which was recently identified as a new INT subunit^45–49^ was present in both interactomes suggesting that it closely interacts with the INTS10-13-14 module. Importantly, we observed many additional factors that are tightly associated to transcription and its regulation in either of the two datasets. Numerous sequence-specific transcription factors (TFs) or co-regulators, subunits of RNAP2 (POLR2A-D/H), elongation factors (ICE1/2, CTR9, PAF1, TCEB2), subunits of chromatin remodelers (INO80, NuRD), and factors involved in histone modifications (ANP32A/B, GRWD1, WDR5) were enriched. Four TFs (ZNF608/609/655/687) and subunit ICE2 of the LEC were copurified with INTS10 and INTS13 (Figure 1C), indicating that they are strongly bound by the module. Interestingly, many of the co-purified TFs have in the past been described as transcription repressors (e.g. ATN1/RERE, GATAD2A, TLE3, ZEB1, ZEB2, ZMYND8, ZNF608, ZNF609), which would be congruent with the attenuation function of INT. These results led us to hypothesize that TFs and other transcription regulators could potentially mediate gene-specific transcription regulation by INT.

**Figure 1.**
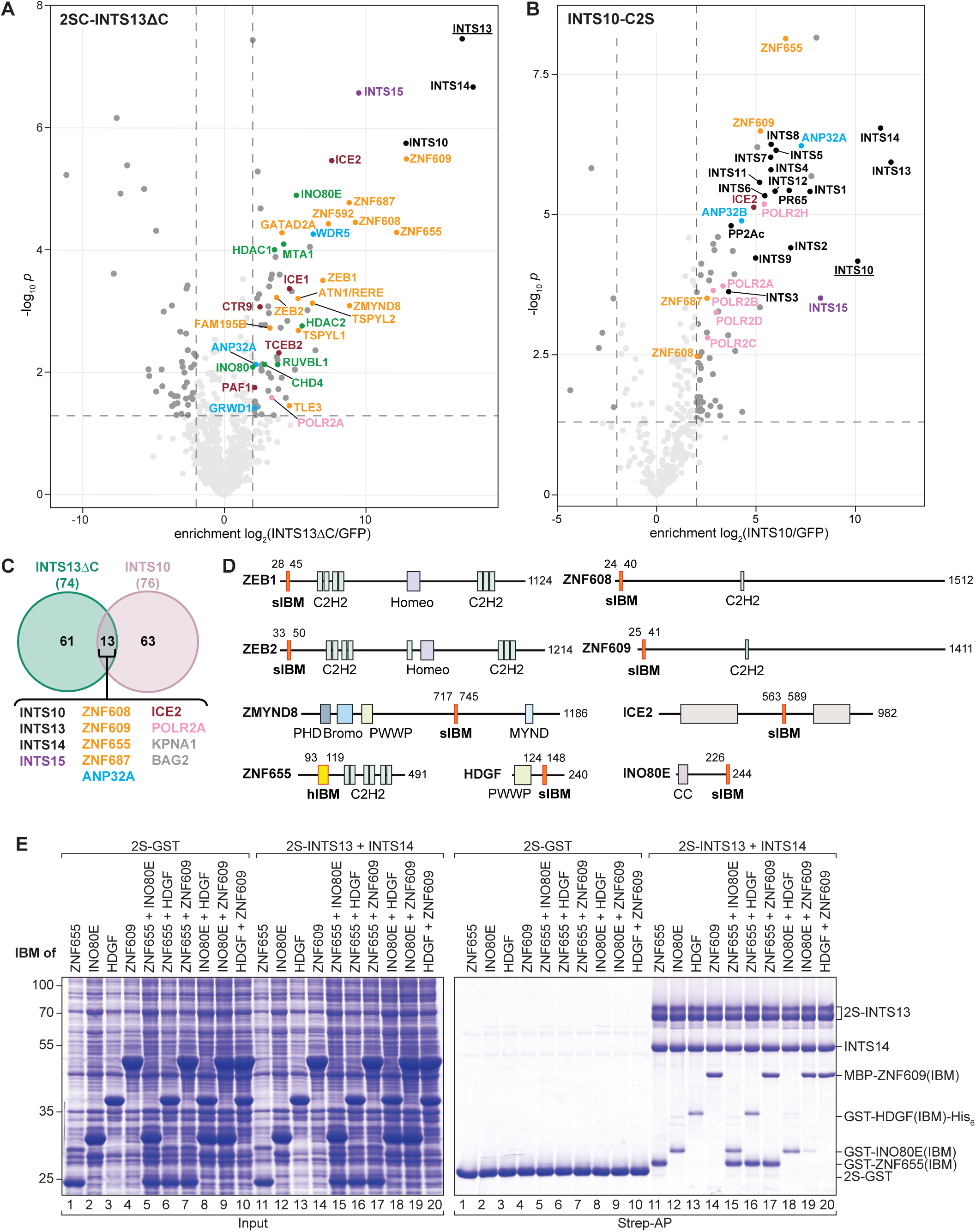
INT binds sequence-specific TFs, chromatin remodelers and little elongation complex via INTS10-13-14. (A, B) Volcano plots displaying proteins identified by TAP-MS of 2SC-INTS13ΔC or INTS10-C2S over GFP control. Significantly enriched proteins (log_2_(INTS13ΔC/GFP > 2, *p* < 0.05) are highlighted (black: INTS, purple: INTS15, orange: TFs, green: chromatin remodelers, pink: RNAP2, red: elongation factors, blue: histone modifiers). (C) Venn diagram showing overlap of significantly enriched proteins in INTS13ΔC and INTS10 TAP-MS datasets with common proteins listed. (D) Domain organization of INTS13-14 interactors highlighting positions of IBM peptides. Domains abbreviations: Bromo – bromodomain, C2H2 – C2H2 zinc finger, CC – coiled coil, Homeo – homeodomain, MYND – myeloid, Nervy, and DEAF-1 type zinc finger, PHD – plant homeodomain, PWWP – Pro-Trp-Trp-Pro domain. (E) Coomassie stained gel of co-APs of MBP or GST-tagged IBMs from *E.coli* extracts with purified 2S-INTS13-14. Single IBMs or two distinct IBMs were added to test for direct interaction or binding competition.

To test whether the detected regulatory factors bind the INTS10-13-14 module directly, we mapped their interacting domains. We screened hits from the INTS13ΔC TAP and performed systematic co-affinity purification (AP) experiments with deletion constructs both after transfection into HEK293T and with recombinant proteins *in vitro* to detect direct binding (Supplemental Figure 1E-H). We identified minimal INT binding motifs (IBM) in several TFs (ZEB1/2, ZMYND8, ZNF608/609/655), as well as in the LEC subunit ICE2 and the INO80 component INO80E that directly contact the INTS13-14 subcomplex (Supplemental Figure 1I-K). These short linear motifs are embedded in long unstructured regions in these otherwise highly diverse factors (Figure 1D). ICE2 as one subunit of LEC is needed for transcription of UsnRNAs.^50,51^ INO80E is a mostly unstructured metazoan-specific subunit of the chromatin remodeling complex INO80.^52^ ZEB1 and ZEB2 are homologous homeobox containing zinc finger (ZF) TFs that are required for embryogenesis and also involved in the epithelial-mesenchymal transition.^53^ ZMYND8 contains a bromo domain, as well as PWWP and ZF DNA binding domains. It acts as a transcription repressor and is involved in DNA damage response.^54^ ZNF608 and ZNF609 are highly homologous (56% similar) proteins with one C2H2 ZF. In mouse embryonic brains they are mutually exclusively expressed and are involved in neurodevelopment.^36^ ZNF655, a poorly characterized mammalian protein containing six C2H2 ZFs, was shown to interact with the proto-oncogene Vav and to be differentially expressed during cell cycle.^55^

Consistent with our data, previously published interactomes of ZMYND8, ZEB1, LEC subunit ICE1, and mouse ZNF609, contain several INTS in addition to INTS10-13-14,^36,51,56–58^ and conversely numerous INTS interactomes also feature ZMYND8, ZNF609, and ZNF655.^35,37,47,59–63^ Together these data indicate that the regulatory factors interact with the whole INT by directly binding the INTS10-13-14 module and that this function is conserved to vertebrates (Supplemental Figure 2B). We therefore propose that INTS10-13-14 form the INT TF binding module (ITFM).

To examine whether, the INT binding partners such as INO80E are embedded in their native functional complexes when they contact INT, we affinity purified INO80 after transfection into HEK293T and indeed detected not only its direct partner INO80E but also INTS13 and INTS1 (Supplemental Figure 2A).

### INT is bound by two different types of motifs

Next, we tested whether different IBMs compete for INTS13-14, and found that only the motif of ZNF655 is compatible with simultaneous binding of other IBMs (Figure 1E), indicating that it contacts a non-overlapping surface of INTS13-14. Indeed, sequence alignments and secondary structure predictions of IBMs from all binders (Figure 2A, I, and Supplemental Figure B) suggest two different classes of motifs. The first class, which we termed α-helical (h)IBM, is only present in ZNF655. BLAST searches in the human proteome with hIBM did not reveal any other proteins with this motif. In contrast, all other direct binders (ICE2, INO80E, ZEB1/2, ZMYND8, ZNF608/609) have in common a short β-strand (LxID) embedded in a negatively charged environment, which we therefore termed strand (s)IBM. The high variation and very short conserved section of the sIBM complicate detection of this motif in other proteins proteome-wide. However, targeted sequence inspection of the other interactors of INTS13 revealed a putative sIBM in TLE3 (transducin-like enhancer protein) that is additionally maintained in other TLE paralogs (TLE1-4, Supplemental Figure 2C), but we could not detect binding in pulldown assays with recombinant proteins (Supplemental Figure 1I). Several other detected interactors of INTS13ΔC are probably binding indirectly. TSPYL2 (testis-specific Y-encoded-like protein 2), ZNF592 and ZNF687 for example are likely associated via ZMYND8 based on our co-purification experiments and published MS data (Supplemental Figure 2D).^58,62,64^

**Figure 2.**
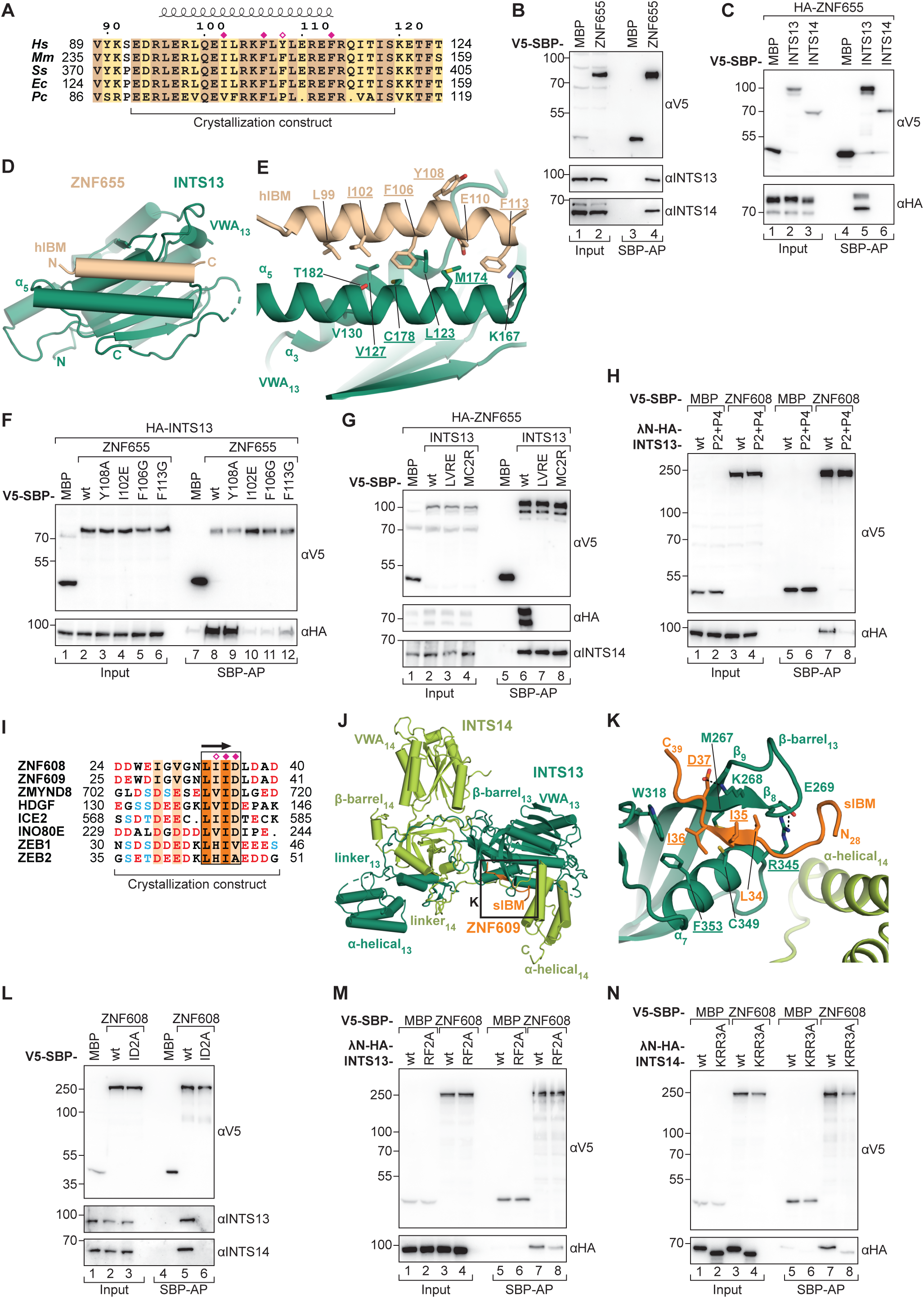
Two surfaces on INTS13-14 serve as binding platforms for distinct types of motifs. (A) Sequence alignment of ZNF655 hIBM in orthologs from *Homo sapiens* (*Hs*), *Mus musculus* (*Mm*), *Sus scrofa* (*Ss*), *Equus caballus* (*Ec*), *Procavia capensis (Pc*). Residues with complete conservation (dark yellow) and 70% similarity (light yellow) are highlighted. α-helix as observed in the INTS13-ZNF655 structure is denoted above the alignment. Red diamonds indicate residues mutated in this study and filled diamonds mark interface residues. (B, C, F-H, L-N) Western blots of inputs and elutions of co-APs from HEK293T with V5-SBP-tagged proteins as baits. (B) Co-AP of endogenous INTS13-14 with V5-SBP-ZNF655. (C) Co-AP of λN-HA-ZNF655 with V5-SBP-INTS13 or V5-SBP-INTS14. (D, E) Crystal structure of complex between INTS13 VWA domain (green) and ZNF655 hIBM (wheat). (E) Side chains of interface and mutated residues are shown as sticks. Mutated residues are underlined. (F) Co-AP of λN-HA-INTS13 with wt or mutant V5-SBP-ZNF655. (G) Co-AP of λN-HA-ZNF655 with wt or mutant V5-SBP-INTS13. (H) Co-AP of wt or mutant λN-HA-INTS13 with V5-SBP-ZNF608. (I) Sequence alignment of sIBMs from INTS13-14 interactors. Residues with complete conservation (orange) and 70% similarity (pale orange) are highlighted. The black box indicates the conserved LxID motif. The β-strand as observed in the INTS13-14-ZNF609 structure is denoted. Red and filled diamonds indicate mutated and interface amino acids, respectively. Negatively charged or phosphorylated residues are highlighted in red or blue. (J, K) Crystal structure of INTS13-14 (dark and light green) in complex with ZNF609 sIBM (orange). (K) Sidechains of interface residues are shown as sticks, mutated residues are underlined. Dotted lines represent hydrogen bonds. (L) Co-APs of endogenous INTS13-14 with wt or mutant V5-SBP-ZNF608. (M) Co-APs of wt or mutant λN-HA-INTS13 with V5-SBP-ZNF608. (N) Co-APs of wt or mutant λN-HA-INTS14 with V5-SBP-ZNF608. Removal of three positively charged residues in λN-HA-INTS14(KRR3A) leads to faster migration in SDS-PAGE compared to wt.

### The ITFM displays two conserved contact sites for TFs

In order to reveal the molecular basis of INT binding by the two motif types, we solved their structures in complex with INTS13-14 (Figure 2). ZNF655 co-precipitates endogenous INTS13-14 from HEK293T but only interacts with V5-streptavidin binding peptide (SBP)-tagged INTS13 not V5-SBP-INTS14 upon co-transfection (Figure 2B, C). Since N-terminal tags on INTS14 hinder its binding to INTS13, this observation indicated direct interaction of ZNF655 with INTS13. Indeed, using APs with recombinant single domain constructs of INTS13 revealed that its VWA domain is sufficient for binding ZNF655 hIBM (Supplemental Figure 2E). We crystallized a minimal complex between ZNF655 hIBM (aa 93-119) and the INTS13 VWA domain (aa 1-256Δ27-48, with deletion of a long surface loop to facilitate crystallization) and solved its structure at 3.2 Å using molecular replacement (Figure 2D and Supplemental Figure 2F-I). The amphipathic helix of ZNF655 hIBM is well ordered (Supplemental Figure 2G) and binds to an accessible and conserved hydrophobic surface on the INTS13 VWA domain that is formed by α-helices 3 and 5 (Figure 2E and Supplemental Figure 2H). Interface mutations in V5-SBP-tagged ZNF655 (I102E, F106G, F113G) strongly impaired co-purification of HA-tagged INTS13 from HEK293T compared to ZNF655 wild type (wt), while removal of a residue that faces away from the interface (Y108A) had no effect (Figure 2F). Analogously, mutations in the hydrophobic cleft of V5-SBP-INTS13 (L123R, V127E [LVRE]; M174R, C178R [MC2R]) completely abolished interaction with HA-ZNF655 without affecting INTS14 binding (Figure 2G). Thus, ZNF655 binds INT on INTS13’s VWA domain via the conserved hIBM also in cells.

For structural characterization of the sIBM-INTS13-14 complex, we chose the sIBM of ZNF609 because it displayed the highest affinity in competition assays (Figure 1E). Interaction of sIBMs containing proteins with INTS13 is abolished by mutations that disrupt INTS13-14 complex formation in co-APs from HEK293T (P2+P4),^44^ suggesting that they contact a composite surface (Figure 2H, Supplemental Figure 3A). We mapped interaction to a minimal complex comprising INTS13 VWA-β-barrel and INTS14 α-helical domain (data not shown). Because this surface is close to the C-terminus of INTS14, we attempted to stabilize sIBM binding, which was sensitive to high salt, by increasing its local concentration, and fused it to the INTS14 C-terminus via a flexible serine-glycine linker (Supplemental Figure 3B). The INTS13-14-ZNF609 sIBM complex was crystallized and resolved to a resolution of 2.5 Å using molecular replacement (Figure 2J and Supplemental Figure 3C-G). Additional unassigned electron density was observed between INTS13 β-barrel and INTS14 α-helical domain that nicely accommodates ZNF609 sIBM (Supplemental Figure 3C). The conserved LxID motif of ZNF609 forms a short β-strand as predicted and binds antiparallel to strand β8 of INTS13. As a consequence of sIBM binding, strand β8 of INTS13 β-barrel is extended and the adjacent loop connecting to strand β9 (M267-Y279) becomes ordered (Figure 2K). Together, β8 and helix α7 of INTS13 create a conserved hydrophobic pocket into which L34 and I36 of ZNF609 are protruding. In addition, D37 of ZNF609 forms a hydrogen bond with K268 of INTS13. Finally, the negatively charged residues located N-terminally of the LxID motif are positioned towards the α-helical domain of INTS14, which exhibits an overall positive surface charge near the peptide (Supplemental Figure 3E, F). To further confirm that the structure accurately reflects binding between ZNF609 and INTS13-14 in solution, as well as general sIBM-INT binding in cells, we tested the effect of structure-based mutations in co-APs. As expected, binding of recombinant ZNF609 sIBM to purified INTS13-14 complexes in *trans* was abrogated by removing the Ile sidechain (I36A) that points into the hydrophobic surface pocket on INTS13, while equivalent mutation of the neighboring Ile that faces away from the interface (I35A) has no effect (Supplemental Figure 3H). In HEK293T, removal of sidechains at positions equivalent to ZNF609 I35 and D36 (ID2A or IV2A) in all V5-SBP-tagged sIBM containing full length interactors (ZNF608, ZNYMD8, ZEB1, INO80E, ICE2), completely abolished their interaction with endogenous INTS13-14, as well as with INTS1 or INTS4, where this was tested (ZMYND8, INO80E, ICE2) (Figure 2L, 3A-C, and Supplemental Figure 3I), without abrogating binding to their other known direct partners (Figure 3B and Supplemental Figure 3J). Analogously, disrupting the hydrophobic pocket on the β-barrel domain of V5-SBP-INTS13 (R345A, F353A [RF2A]) did not disrupt formation of the INTS13-14 complex (Supplemental Figure 3K) but weakened binding of HA-tagged sIBM containing interactors (ZNF608, ZMYND8, INO80E, ICE2) in HEK293T, although to different degrees, indicating that for individual sIBM proteins slightly different aspects of the INTS13-14 surface contribute to binding (Figure 2M and 3E-G).

**Figure 3.**
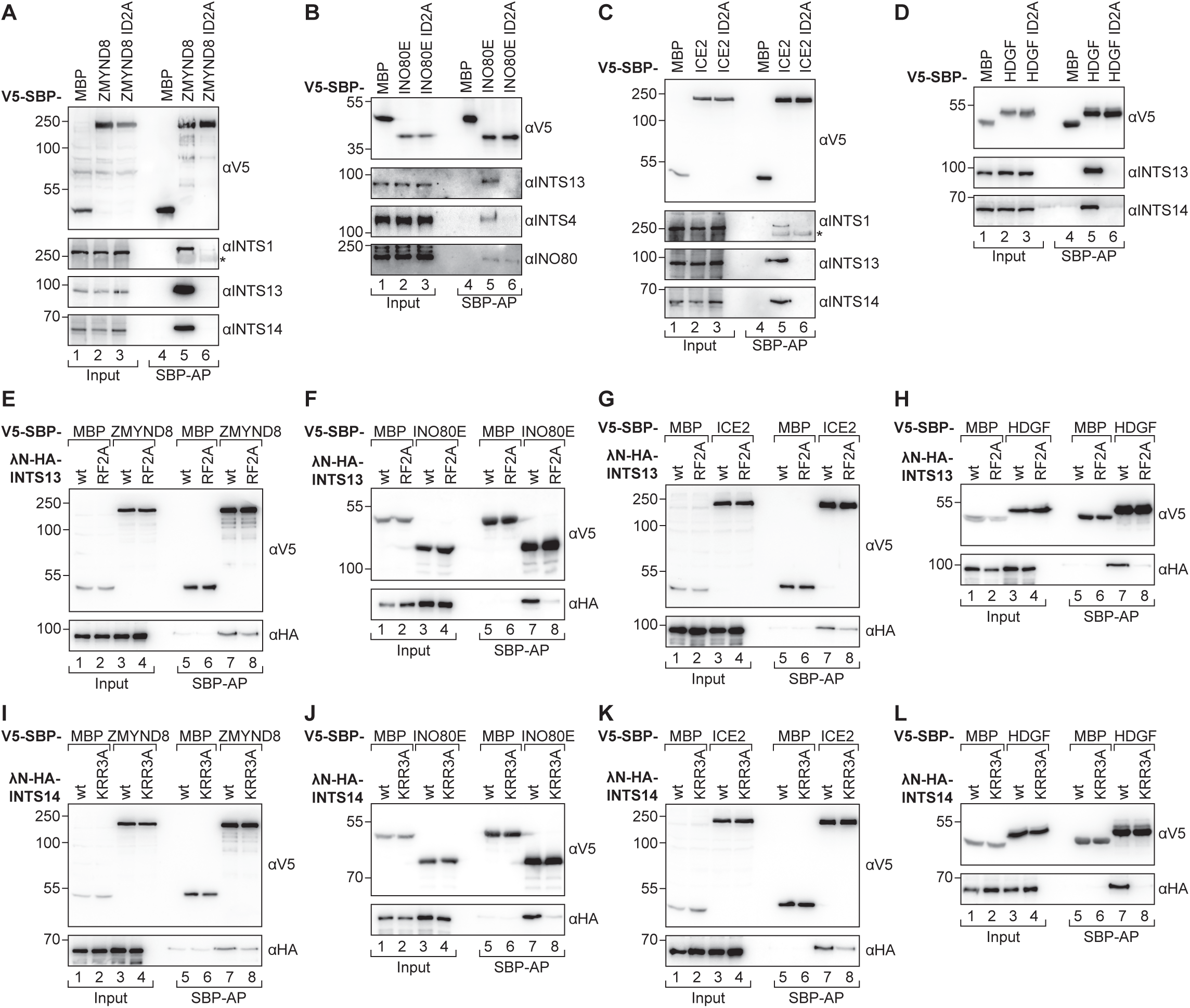
Several TFs, INO80E and ICE2 bind to the same hub on INTS13-14 via related motifs. (A-L) Western blots of inputs and elutions of co-APs from HEK293T cells. (A-D) Co-APs of endogenous proteins with wt or mutant V5-SBP-tagged ZMYND8 (A), INO80E (B), ICE2 (C) or HDGF (D). Asterisks mark unspecific bands. (E-H) Co-APs of λN-HA-INTS13 wt or mutant with V5-SBP-tagged ZMYND8 (E), INO80E (F), ICE2 (G) or HDGF (H). (I-L) Co-APs of λN-HA-INTS14 wt or mutant with V5-SBP-tagged ZMYND8 (I), INO80E (J), ICE2 (K) or HDGF (L).

Given that sIBM binding is sensitive to INTS13-14 complex disruption in cells and high salt concentrations *in vitro*, and that the negatively charged N-terminal sIBM region is positioned towards a positive patch on INTS14, we reasoned that they might be involved in stabilizing sIBM contacts. Indeed, removal of three positively charged side chains from this conserved patch in V5-SBP-INTS14 (K435A, R439A, R442A [KRR3A]) impaired co-AP of all sIBM binders tested (ZNF608, ZMYND, INO80E, ICE2, Figure 2N and 3I-K), without negatively impacting INTS13-14 complex formation which rules out indirect effects (Supplemental Figure 3L). This observation prompted us to test whether the sIBM interaction could be sensitive to presence of RNA, which was previously shown to bind INTS10-13-14.^44,65^ However, we observed no changes to complex stability upon addition of poly-U RNA (Supplemental Figure 3M), suggesting that in cells also during transcription TF-INT binding would not be hampered by presence of RNA.

Taken together, our data show that sIBM-containing proteins bind INT in cells on a conserved composite INTS13-14 interface that consists both of a hydrophobic pocket for a short β-strand and a positive surface patch for contacts with negatively charged motif sections.

### Stress conditions modulate the interactome of INT

INT-mediated gene regulation is required for inducible transcriptional programs. If TFs are responsible for INT gene targeting as a result of different stimuli, this might be reflected in INT’s interactome. Therefore, we repeated TAP-MS of 2SC-INTS13ΔC upon three distinct perturbations that have previously been reported to elicit an INT-dependent response (Figure 4A-C and Supplemental Figure 4A-C): heat shock (42°C) that leads to the induction of chaperones,^32^ glucose starvation that triggers formation of primary cilia,^37^ and epidermal growth factor (EGF) addition after serum starvation that induces immediate early genes.^26,32^ Indeed, we observe changes in the interactome of INTS13ΔC under these three stress conditions in comparison to normal conditions and also compared to each other (Figure 4D). In particular, several TFs are differentially enriched. Most prominently, in all stress conditions HDGF (hepatoma-derived growth factor) is a new significant hit, TLE1 and TLE3 become more strongly enriched, while TSPYL1 is lost.

**Figure 4.**
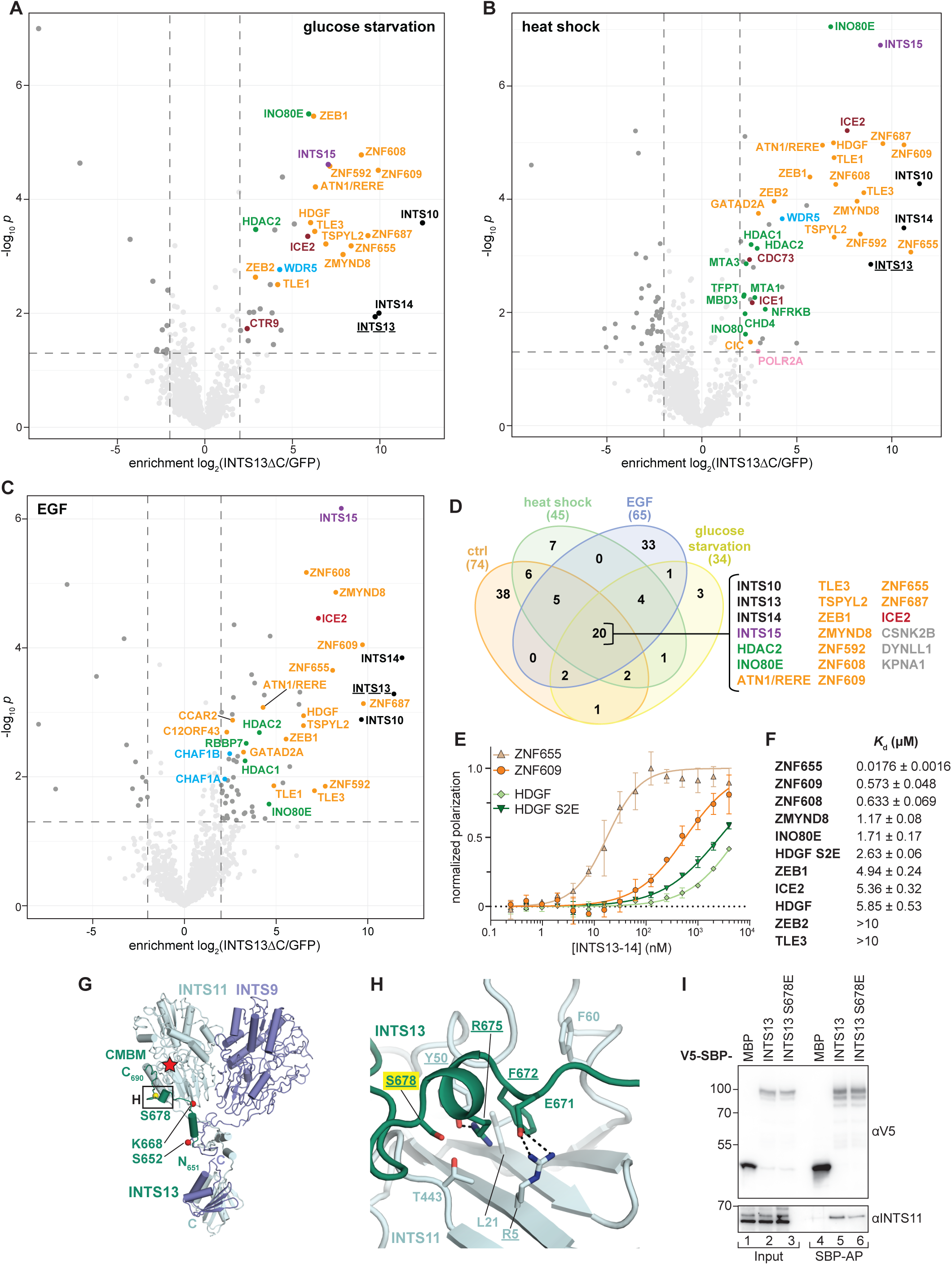
The TF interactome of INT is modulated by stress conditions and could be regulated by PTMs. (A-C) Volcano plots displaying enrichment of proteins identified by TAP-MS of INTS13ΔC over GFP control after 24 h glucose starvation (A), 2 h of 42 °C heat shock (B) or 1 h EGF treatment after 48 h serum starvation (C). Significantly enriched proteins (log_2_(INTS13ΔC/GFP) > 2, *p* < 0.05) are highlighted and colored as in Figure 1. (D) Venn diagram showing the overlap between significantly enriched proteins of INTS13ΔC TAP-MS experiments performed under normal and stress conditions, with commonly identified proteins listed. (E, F) Affinity measurements by fluorescence polarization for GFP-tagged IBMs and INTS13-14. Exemplary binding curves are shown in (E), all other curves are found in Supplemental Figure 4. Error bars indicate standard deviations of means from three experiments. All determined dissociation constants are listed (F) with standard error of means derived from fitting binding curves. (G, H) Structure of INTS9-11 (light, dark blue) bound by INTS13 CMBM (green) generated by reanalysis of published INT cryoEM density (EMD-30473-2). The red star marks INTS11’s active site. Red spheres indicate INTS13 residues mutated in human ciliopathy patients, the yellow sphere marks a phosphorylation site. Sidechains of interface residues are shown and labeled in (H). Dotted lines represent hydrogen bonds. Residues mutated in this study are underlined. The phosphorylation site is highlighted in yellow. (I) Western blot of co-AP of endogenous INTS11 with V5-SBP-INTS13 wt or phospho-serine-mimetic mutant S678E from HEK293T.

HDGF is an endogenous nucleus-targeted mitogen with a PWWP DNA binding domain (Figure 1D) and reported transcription repressor activity.^66^ We subjected recombinant HDGF deletion constructs to co-APs with purified INTS13-14 and mapped direct binding to its middle section (Supplemental Figure 4D). Sequence inspection revealed presence of an sIBM in this region (Figure 2I and Supplemental Figure 2B). Consistently, HDGF sIBM behaves analogous to the other identified sIBMs: It competes with other sIBMs for binding to INTS13-14 (Figure 1E), and its binding to cellular INTS13-14 is disrupted by the same type of mutations that also impair the other sIBM motifs (Figure 3D, H, L and Supplemental Figure 4E).

### Post-translational modifications (PTMs) could modulate TF-INT affinity

To quantify IBM affinities for INTS13-14, we carried out fluorescence polarization assays (FP) with GFP-tagged IBMs (Figure 4E, F and Supplemental Figure 4F). Dissociation constants (*K*_d_) of IBMs from different proteins differed by roughly three orders of magnitude. Consistent with the larger hydrophobic interface of ZNF655 hIBM, it displays the highest affinity for INTS13-14 (*K*_d_=17.6 nM), while most sIBM affinities were ∼30-300-fold lower. As suggested by the competition assays, sIBMs of ZNF608/609 had the highest affinities amongst this class of binders (*K*_d_=0.633 and 0.573 µM), while ZEB2 was the weakest interactor that we tested (*K*_d_ > 10 µM). Although we had not been able to detect TLE3 binding in co-APs, we did observe weak binding by FP (*K*_d_>10 µM).

Further analysis of all sIBM sequences showed that most of the short β-strands are preceded by conserved serine/threonine/tyrosine residues, which are phosphorylated in cells (Figure 2I and Supplemental Figure 2B, C).^67,68^ Given the relevance of the positive patch on INTS14 for sIBM binding in cells, modification of these residues could potentially be exploited by cells to modulate binding affinity of the respective effector for INT. We tested this hypothesis on HDGF sIBM, which has two serines (S132/133) that are modified in cells, because it has one of the lowest affinities of all tested sIBMs and was only significantly enriched under stress conditions. Indeed, replacing both serines with phospho-serine-mimetic glutamate residues (S2E mutant, Figure 4E, F) increased HDGF sIBM affinity for INTS13-14 more than twofold (*K*_d_=2.63 µM (S2E) *vs.* 5.85 µM (wt)). This result indicates that by placing or removing PTMs on the sIBMs, cells could modulate TF-INT affinity in a fast and responsive manner to trigger a quick transcriptional response.

### ITFM incorporation into INT via cleavage module contacts could be modulated by PTMs

INTS13 binds the ICM directly via a short α-helical section at its C-terminus, however the molecular basis of this interaction is not known.^37,44^ Re-inspection of the cryogenic-electron microscopy (cryoEM) density of a published INT structure^30^ revealed unmodeled additional density on the ICM surface (INTS9-11, Supplemental Figure 4G) that was consistent with an AlphaFold2 (AF2) model of CMBM in complex with INTS9-11 (Supplemental Figure 4H). We adjusted the CMBM model to fit the density, which allowed us to place INTS13 aa651-690 confidently (Figure 4G, H and Supplemental Figure 4G, I, J). The INTS13 CMBM stretches along a conserved INTS9-11 surface from CTDs to catalytic domain forming two short α-helices. Mutation of predicted interacting residues in both INTS11 and INTS13 abrogated binding, thereby verifying the CMBM-INTS9-11 model (Supplemental Figure 4K, L).

The structure explains the deleterious effects of human ciliopathy mutations in the INTS13 CMBM.^37^ They likely disrupt CMBM binding by either extending the first α-helix leading to a steric clash (S652L), or by effectively removing two thirds of the interaction surface (K668-frameshift) (Supplemental Figure 4I). Importantly, INTS13 CMBM also contains a documented phosphorylation site on S678 that is involved in the interface with INTS11 (Figure 4G, H),^67^ which could modify its binding affinity. Mutation S678E as a proxy for phosphorylation, indeed weakened binding of INTS13 to endogenous INTS11 in HEK293T (Figure 4I). Together, these data suggest that by decreasing INTS13 affinity for INTS11 via PTMs, cells could modulate incorporation efficiency of ITFM into functional INT and thereby as a consequence also titrate INT-TF recruitment.

### INT and its TF interactors co-regulate stress response genes

To determine whether INT and its TF interactors might co-regulate transcription, we carried out RNA-sequencing (RNAseq) after depletion of either INTS13, HDGF, TSPYL2, ZEB1, ZMYND8, ZNF608, ZNF609, ZNF655, or ZNF687 using siPOOLs (30 different siRNAs pooled, Supplemental Figure 5A,B). Differentially expressed genes (DEGs) upon INTS13 depletion (Figure 5A) were enriched for Gene ontology (GO) terms for various stress responses, including starvation, radiation, oxygen and heat stress, consistent with INT regulation of inducible genes (Figure 5B). Another GO-term points to involvement in neurogenesis, which is in line with neurodevelopmental disorders connected to INT mutations.^37,43^ As expected for an attenuator ∼2/3 of all DEGs (552 of 863) were upregulated upon depletion. Given that the DEGs include mRNAs of TFs (GO-term “double-stranded DNA binding”) the downregulated DEGs (311) may be due to secondary effects as suggested before.^69^

**Figure 5.**
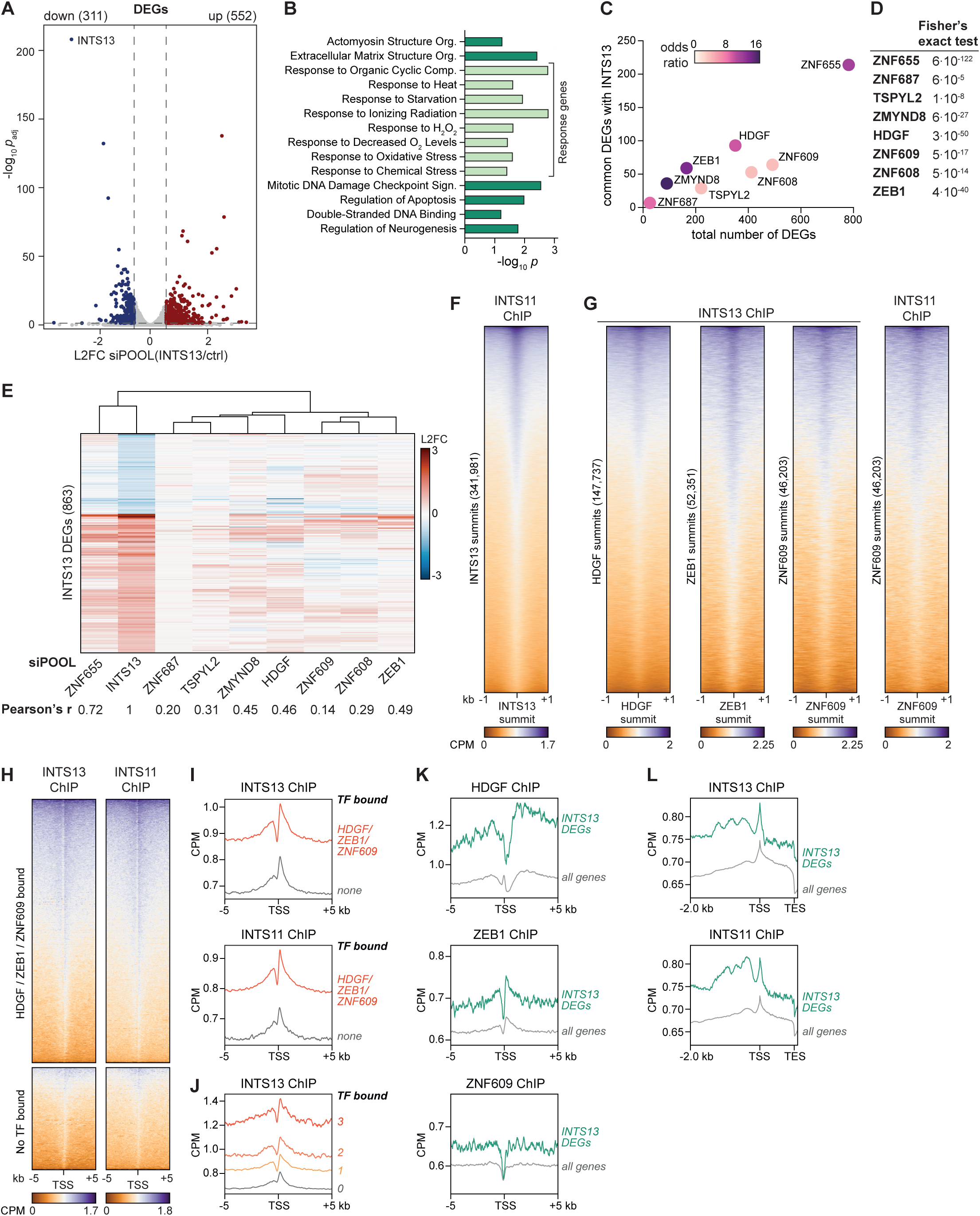
INT binding TFs co-regulate overlapping gene sets with INT and co-localize on chromatin with INT. (A) Volcano plot of RNAseq data showing L2FCs and *p*_adj_ of transcripts from HEK293T treated with INTS13 siPOOL relative to ctrl siPOOL (n=2). Significantly (|L2FC| > 1.5, *p*_adj_ < 0.05) down-(blue) and upregulated (red) DEGs are colored, INTS13 mRNA is labelled. (B) GO-term analysis of INTS13 DEGs, showing significantly enriched terms (*p* < 0.05) ordered by decreasing number of associated genes. (C) Scatter plot showing total number of DEGs in each TF depletion versus number of DEGs in common with INTS13 depletion. Colors of data points represent odds ratio of DEG overlap for each TF depletion with INTS13 depletion. (D) Fisher’s exact test *p*-values for significance of overlap between DEGs in TF and INTS13 depletions. (E) Heatmap showing L2FC of DEGs identified upon INTS13 depletion in each TF knockdown RNAseq sample. Correlation with INTS13 DEGs is listed below as Pearson’s *r*-value. (F, G) Heatmaps of ChIPseq signals (counts per million reads, CPM) of INTS11 around INTS13 peak summits (F), of INTS13 and INTS11 around HDGF, ZEB1 and ZNF609 peak summits (G). (H) Heatmaps of INTS13 and INTS11 ChIPseq signals around TSSs grouped into genes bound by minimally one TF (HDGF, ZEB1 or ZNF609) or none. (I) Profiles of INTS13 and INTS11 ChIPseq signals around TSSs on genes bound by minimally one TF (orange) or none (grey). (J) Profiles of INT13 ChIPseq signal around TSSs of genes bound by 0 (grey), 1 (yellow), 2 (light orange), or 3 (dark orange) TFs. (K) Profiles of HDGF, ZEB1 and ZNF609 ChIPseq signals around TSSs on INTS13 DEGs (green) and all genes (grey). (L) Profiles of INTS13 and INTS11 ChIPseq signals between TSS and TES as well as 2 kb upstream of TSSs at INTS13 DEGs (green) and all genes (grey).

Depletion of interacting TFs also led to considerable misregulation of genes, which significantly overlapped with DEGs upon INTS13 depletion as judged by Fisher’s exact test (Figure 5C, D). More than 40% of all INTS13 DEGs (356 transcripts) are also significantly misregulated upon one of the TF depletions. Of all TFs knockdowns, ZNF655 depletion displays the most striking similarity to the DEG pattern after INTS13 depletion (Figure 5E, Pearson’s r=0.72), especially for the up-regulated genes, which correlates with the high affinity of ZNF655 for INTS13. Furthermore, there is notable diversity in the commonly misregulated genes after each TF and INTS13 knockdown (with the exception of paralogs ZNF608/609), which is highlighted by the diverse biological pathways that are enriched among DEGs for each TF knockdown (Supplemental Figure 5C). All of the stress response pathways (apart from H_2_O_2_) affected by INTS13 depletion are recapitulated by at least one of the TFs. Taken together, our data suggest that INT binds different TFs to co-regulate distinct, TF-specific biological pathways, which are often, but not limited to, stress response.

### INT subunits co-localize with TFs on the genome

Based on the strong functional interaction between INTS13 and its TF interactors, we hypothesized that these TFs may recruit INT to specific genomic loci. Therefore, we performed chromatin immunoprecipitation followed by sequencing (ChIPseq) from HEK293T using antibodies against INT subunits from two independent modules (INTS13 of ITFM, INTS11 of ICM), and of some TFs for which reliable antibodies are available (HDGF, ZEB1, ZNF609). Both INTS11 and INTS13 enrich immediately downstream and upstream of TSSs, which is consistent with INT’s previously described roles at promoter-proximal pause sites^18^ and in restricting antisense transcription^20–22^ (Supplemental Figure 5D). Although, INTS11 is clearly accumulated at INTS13 peaks (Figure 5F), it is only present on ∼55% of all INTS13-bound genes (Supplemental Figure 5E) consistent with a previous study.^35^ In addition to TSSs, both INTS ChIP peaks are located to a large proportion (73% INTS13, 43% INTS11) at promoters (5 kB upstream of TSS), which would be consistent with their potential recruitment to these loci by TFs (Supplemental Figure 5F). In support of this model, both INTS ChIP signals around TSSs are increased at INTS13 promoter-bound genes, relative to non-promoter bound genes (Supplemental Figure 5G, H). While, almost all INTS11 promoter-bound genes also display INTS13 at their promoter, ∼2/3 of INTS13 promoter-bound genes do not recruit INTS11 (Supplemental Figure 5I), suggesting that ITFM can attach to promoters independently of the rest of INT. This observation is consistent with the higher cellular abundance of INTS13-14 compared to other INTS^62,70^ and could indicate that incorporation of ITFM into INT via INTS13-CMBM might indeed be regulated (Figure 4G-I).

In keeping with direct TF-recruitment of INT, INTS11/13 both co-localize with HDGF, ZEB1 and ZNF609 summits (Figure 5G and Supplemental Figure 5J). Their ChIP signal is more broadly dispersed upstream and downstream of the TSS at genes that are bound by HDGF, ZEB1, or ZNF609, compared to genes that do not display peaks of these TFs (Figure 5H, I). Target genes of the three TFs also display significantly higher occupancy of both INTS around TSSs, relative to genes not bound by any of these TFs, and the increase in ChIP signal scales with how many of the TFs are present at the respective genes (Figure 5I, J). Although these TFs all bind to the same site on ITFM, this observation suggests that a higher local concentration of redundant INT binders on chromatin leads to a concomitant enrichment of INT. It is important to note that of 17,695 INTS13 promoter-bound genes, 4533 are not bound by any of the TFs ChIP-ed in this study (Supplemental Figure 5K), suggesting that INT may be recruited to the remaining promoter-bound genes through interactions with other TFs, such as ZNF655.

Consistent with the notion that TF-promoter recruitment could explain INT-mediated transcription regulation, we observe that ∼2/3 of INTS13 DEGs are bound by INTS13 at their promoters, whereas only 99 show INTS13 peaks in other regions (Supplemental Figure 5L, M). Furthermore, HDGF, ZEB1, ZNF609, and INTS11/13 ChIP signals are higher around TSSs of INTS13 DEGs (Figure 5K, L). Strikingly, INTS13 DEGs display a unique enrichment of INTS13 and INTS11 peaks in a 2 kB region upstream of the TSS, which is not observed at other genes (Figure 5L) and could reflect TF-INT complexes at promoters. Taken together, our data indicate that INT is recruited by TFs to promoters of its target genes to elicit changes in their expression.

### INTS13 cooperates with HDGF and ZMYND8 to modulate glucose stress response

Due to the reported involvement of INT in regulation of inducible genes and the enrichment of stress response genes in DEGs of several of its direct TF binders, we sought to examine to which extent they coordinate the transcriptional response to stress using glucose starvation as a model. We performed RNAseq after 24 h glucose deprivation of HEK293T treated with ctrl siPOOL or depleted of INTS13, HDGF or ZMYND8 (Supplemental Figure 6A-E), because these TFs regulate starvation genes (Supplemental Figure 5C). We defined “glucose starvation response genes” as DEGs in starved *vs*. unstarved cells under ctrl depletion conditions (1009 genes, Figure 6A and Supplemental Figure 6D). To focus on transcripts that are regulated by INTS13, we separated genes that are still differentially expressed after starvation when INTS13 is absent, leaving 873 INTS13-dependent genes (Figure 6A, B and Supplemental Figure 6E). In general, all starvation response genes, both INTS13-dependent and independent, tend to be upregulated upon glucose removal (Figure 6C and Supplemental Figure 6F). However, INTS13-independent genes display significantly higher expression levels than INTS13-dependent genes, indicating that INT regulates genes with lower expression levels as has been suggested before.^20,71^ As a whole, starvation response genes are upregulated by INTS13 depletion only when glucose is present but are insensitive to INTS13-loss when glucose is absent, suggesting an involvement of INTS13 in repressing these genes under normal circumstances (Figure 6D and Supplemental Figure 6G). This effect becomes more pronounced when we focus on INTS13-dependent genes, in contrast to INTS13-independent genes that show no significant misregulation when INTS13 is depleted regardless of growth condition. Attenuation is the best described mechanism for INT-mediated regulation of inducible genes.^17^ To examine how attenuation influences glucose starvation genes, we further classified the group of INTS13-dependent starvation response genes into those that are down- or upregulated by INTS13 depletion in glucose-containing medium (287 *vs.* 586 genes), because the latter are candidates for INT attenuation (Figure 6E, F and Supplemental Figure 6H, I). If glucose removal cancels attenuation of starvation response genes, then these transcripts should only be upregulated by INT inactivation (i.e. INTS13 depletion) when glucose is supplied but not when glucose is absent. In keeping with this model, expression levels of INT attenuated genes increase upon starvation (Figure 6E and Supplemental Figure 6H, “up”) and also upon INTS13 depletion in normal medium (Figure 6F, “up”), but show no expression change after INTS13 depletion under starved conditions. Conversely, for non-INT attenuated transcripts, glucose removal has only slight effects on expression levels (Figure 6F and Supplemental Figure 6I, “down”) and INTS13 depletion leads to overall downregulation no matter whether glucose is present or not (Figure 6F, “down”).

**Figure 6.**
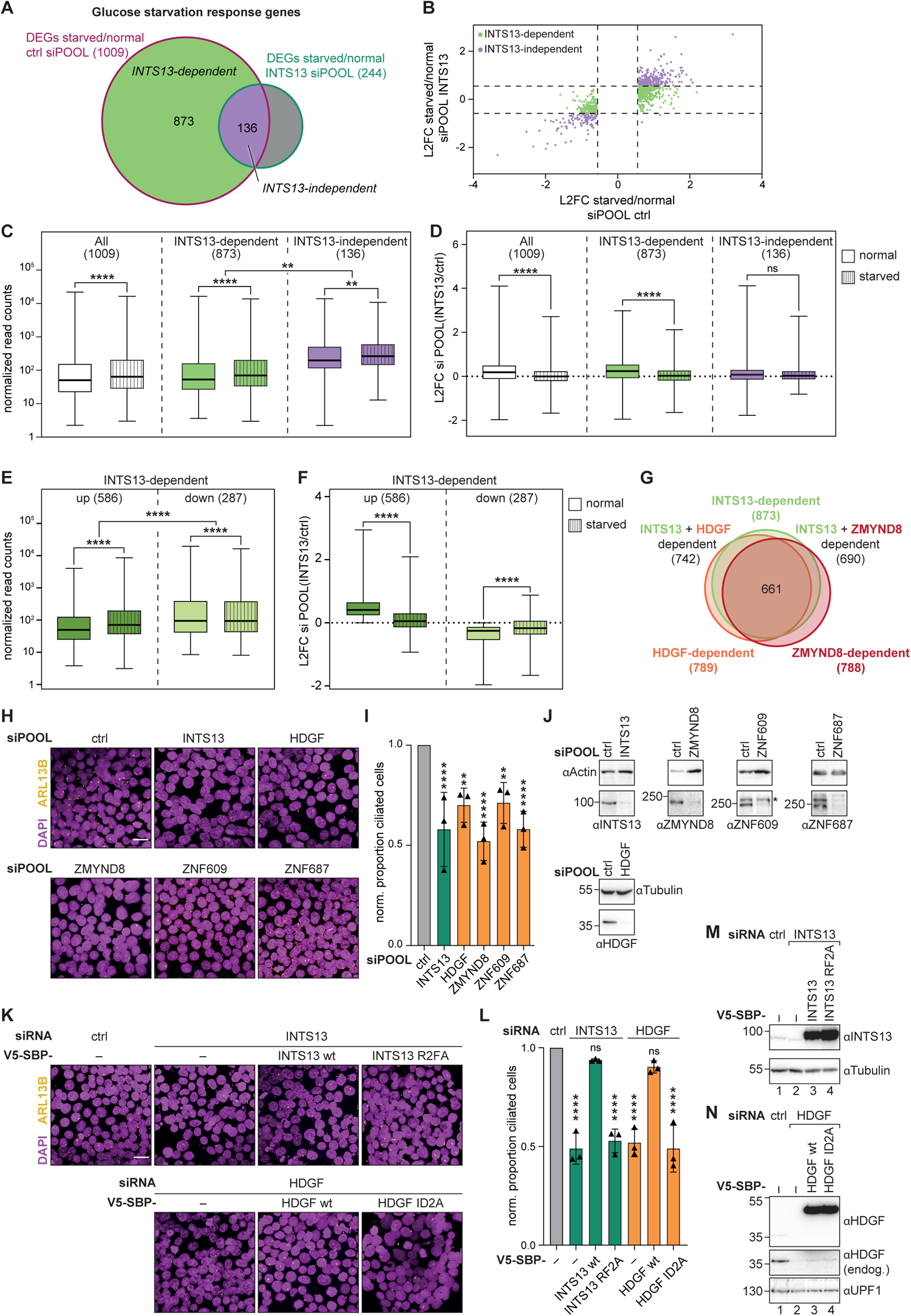
Response to glucose starvation depends on TF-INT binding. (A) Venn diagram of overlap between DEGs upon glucose starvation under ctrl or INTS13 depletion. Glucose starvation DEGs only found upon ctrl depletion are defined as “INTS13-dependent” (green), while those transcripts that are also changed upon starvation when INTS13 is depleted are defined as “INTS13-independent” (purple). (B) Scatter plot showing L2FCs of transcript levels upon glucose starvation in ctrl and INTS13 siPOOL treated cells. INTS13-dependent (green) and independent genes (purple) are colored. (C) Box plot of normalized read counts of all (left, white), INTS13-dependent (middle, green) and independent (right, purple) starvation response genes under normal (plain) and starvation (striped) conditions. (D) Same as (C) but plotting L2FCs of these genes upon INTS13 siPOOL treatment. (E) Box plot of normalized read counts of those INTS13-dependent glucose starvation response genes that are up-(left, dark green) or downregulated (right, light green) upon INTS13 depletion by siPOOL (under normal conditions). Comparison between their read counts under normal (plain) and glucose starvation (striped) conditions is shown. (F) Same as (E) but plotting L2FCs of these genes upon INTS13 siPOOL treatment. (C-F) Statistical significance was calculated using paired t-test within gene sets and unpaired t-test across gene sets. All boxes show 25th–75th percentiles and whiskers depict minimum and maximum values. (G) Venn diagram of the overlap between INTS13-, HDGF- and ZMYND8-dependent glucose starvation response genes. (H) Immunofluorescence images of HEK293T stained for the ciliary axoneme component ARL13B (yellow) and DAPI (magenta) treated with ctrl, INTS13, HDGF, ZMYND8, ZNF687, or ZNF609 siPOOLs. (I) Quantification of number of ciliated cells over total number of nuclei under INTS13 (green) or TF (orange) depletion conditions normalized to ctrl depletion (grey). (J) Representative Western blots of depletions used in (H, I), blots of the other replicates are found in Supplemental Figure 6. Actin or tubulin served as loading control. (K) Immunofluorescence imaging as in (H) but cells were treated with ctrl, INTS13 or HDGF siRNAs and co-transfected with siRNA-resistant V5-SBP tagged wt or mutant INTS13 or HDGF. (L) Quantification of rescue assays analogous to (I). For (I) and (L) N_cells_ > 500 per replicate for three biological replicates. Statistical significance compared to respective ctrl depletion was calculated using ANOVA. (M, N) Representative Western blots of depletions and transfections used in (K, L), blots for the other replicates are found in Supplemental Figure 6. For all panels *p*-values are indicated as ns > 0.05, ** < 0.01, and **** < 0.0001.

To understand how TFs are involved in coordinating the response with INT, we identified HDGF- and ZMYND8-dependent starvation response genes analogous to INTS13 (Supplemental Figure 6J, K). Almost all INTS13-dependent starvation response genes overlap with either HDGF- or ZMYND8-dependent genes with 75% overlapping with both sets (Figure 6G and Supplemental Figure 6L), pointing towards a role for both TFs in INT-regulated starvation response.

### INT-TF binding is required for starvation-induced ciliogenesis

Primary cilia formation is an important cellular response to starvation and other stress in many human cell types.^72^ INT is linked to sensory cilia formation, because INTS13 mutations that disrupt ITFM incorporation into INT cause a human ciliopathy.^37^ Since additionally our above results indicate that INTS13 and several of its TF-binders are required for transcriptional glucose starvation response, we next tested whether TF binding to INT is required for ciliogenesis in HEK293T. We determined the fraction of ciliated cells upon glucose deprivation using immunofluorescence (IF) microscopy after knockdown of INTS13, or TFs that regulate starvation response genes using siPOOLs (HDGF, ZMYND8, ZNF609, ZNF687, Supplemental Figure 5C) and normalized them to ctrl depletion (Figure 6H-J and Supplemental Figure 6M). As expected, INTS13 depletion reduced the normalized proportion of ciliated cells to 58% (Figure 6H, I). Since knockdowns of HDGF (70%), ZMYND8 (52%), ZNF609 (71%) or ZNF687 (58%) similarly impair ciliogenesis, our results altogether suggest that INT cooperates with TFs to orchestrate a cellular response to glucose starvation. To test whether direct binding between INT and the TFs is required for this orchestration, we performed rescue experiments under glucose starvation with binding interface mutants (Figure 6K-N and Supplemental Figure 6N, O). Ciliogenesis defects induced by INTS13 siRNA depletion (49%) were rescued by supplying siRNA-resistant INTS13 wt (94%) but not by the INTS13 RF2A mutant (53%) that cannot bind any of the sIBM containing TFs. Conversely, impaired cilia formation upon HDGF siRNA depletion (52%) could only be recovered by co-transfection of siRNA-resistant HDGF wt (90%) but not by the HDGF ID2A mutant (49%) that is defective for INTS13 binding. Thus, it is crucial for an intact cellular starvation response that INT can bind the TFs directly.

### INTS15 and DSS1 are stable subunits of INT

As a next step we aimed to reveal whether TF binding on INT would be architecturally compatible with recognition of paused RNAP2. None of the published structures of INT, neither in isolation nor bound to PEC, contain all INTS and in particular the whole ITFM is missing.^13,16,30^ Therefore, we used a combination of biochemical mapping, AF2 predictions and reanalysis of published cryoEM maps to further complete the INT structure.

Consistent with several recent studies that identified C7ORF26 as a subunit of INT (INTS15),^45–49^ we detected it consistently in all our INTS13ΔC MS datasets (Figure 1A and 4A-D), indicating that INTS15 directly associates with ITFM. Systematic co-APs with recombinant INT subcomplexes revealed that INTS15 in addition to INTS10-13-14 also binds the core subcomplex INTS5-8 (Figure 7A and Supplemental Figure 7A), and we could further narrow it down to the N-termini of INTS5 and INTS10, which form a stable minimal complex with INTS15 *in vitro* (Supplemental Figure 7B). We generated high confidence AF2 models of this minimal INTS5N-15-10N assembly as well as the INTS10-14 contact (Supplemental Figure 7C-F, I-L) that we had previously identified biochemically to map to the INTS14 MIDAS pocket.^44^ The relevance of the predicted binding interfaces (INTS10-14, INTS10-15, INTS5-15) was verified using co-APs of model-guided mutants in HEK293T (Figure 7B-D and Supplemental Figure 7G, H). While all mutations efficiently disrupted the targeted interactions, they did not affect association of other respective INTS binding partners. Structural superposition of the individual models together with our previously resolved INTS13-14 structure^44^ yielded a model of the entire ITFM with INTS15 forming a bridge to the INT core (Figure 7E), which is consistent with a recent study that came to similar conclusions.^47^

**Figure 7.**
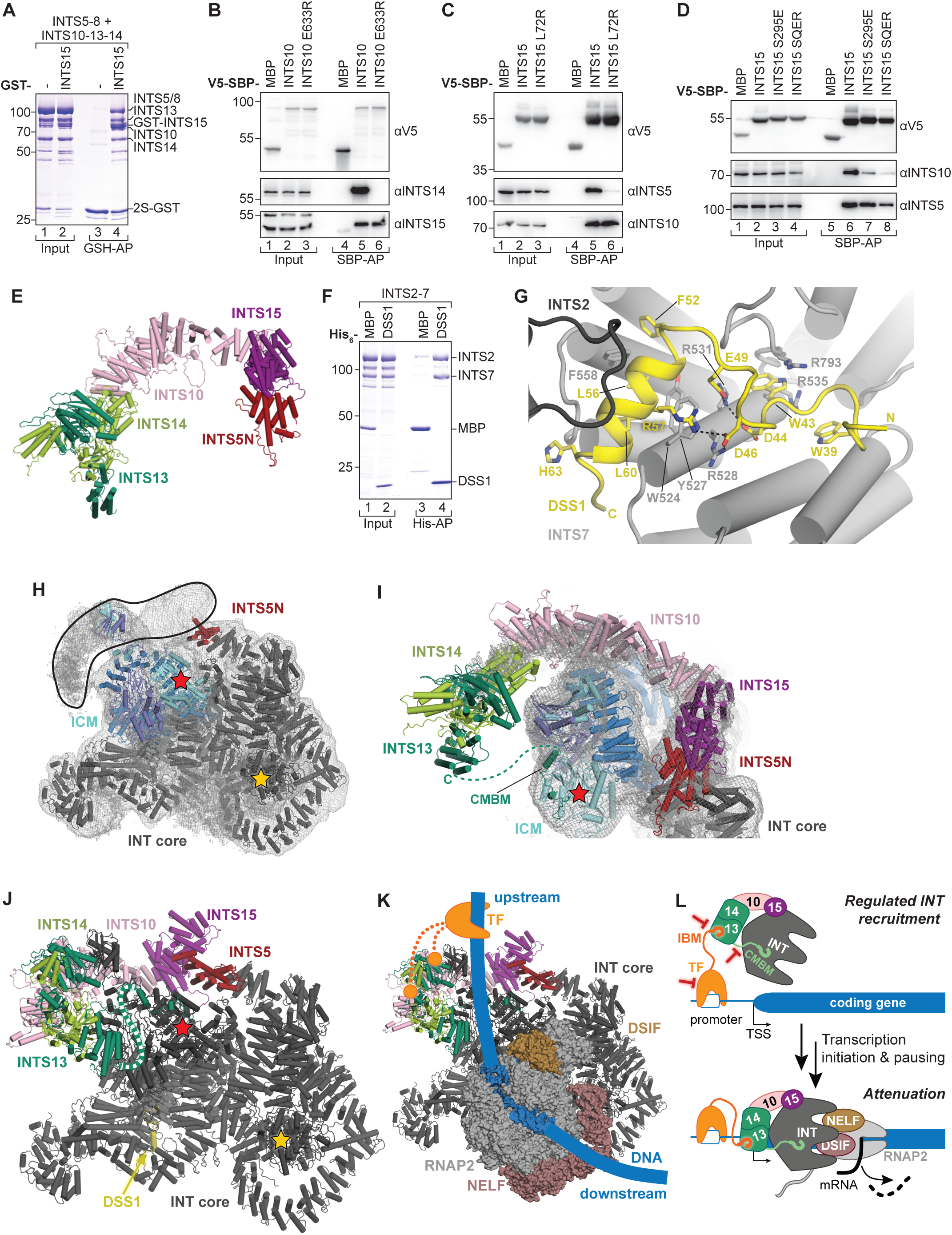
The INTS10-13-14 module is poised for recruitment of INT to promoter-proximally paused RNAP2 via TFs. (A) Coomassie stained SDS-PAGE of co-APs of purified GST-INTS15 with purified INTS5-8 and INTS10-13-14 complexes. (B-D) Western blots of co-APs of endogenous proteins from HEK293T with wt or mutant V5-SBP-tagged INTS10 (B) or INTS15 (C, D). (E) Model of the INTS13-14-10-15-5N complex based on AF2 predictions and the INTS13-14 crystal structure (PDB-ID: 6SN1). (F) Coomassie stained SDS-PAGE of co-APs of purified INTS2-7 complex with purified His_6_-DSS1. (G) Close-up of the interface between INT2-7 (dark and light grey) and DSS1 (yellow). Sidechains of interface residues are shown and labeled. Dotted lines represent hydrogen bonds. (H) CryoEM density (grey mesh) and structure of the published INT complex (PDB ID: 7CUN), consisting of INTS1-2-5-6-7-8-PP2A (INT core, grey), and INTS4-9-11 (ICM, royal, dark and light blue). The INTS5 N-terminus is colored red, red and yellow stars mark INTS11’s and PP2Ac’s active site. Unmodeled density is highlighted by a black outline. (I) Superposition of the INTS13-14-10-15-5N model from (A) onto INTS5N of the published INT structure shows that the additional subunits explain the unmodeled EM density. (J) Completed INT model with added INTS10-13-14-15, DSS1 and INTS13 CMBM to the published structure (PDB ID: 7CUN). The dotted green line indicates the unstructured linker that connects INTS13 α-helical domain and CMBM. The red star marks INTS11’s active site. (K) Superposition of the completed INT model onto the published INT-PEC structure (PDB-ID: 7PKS). RNAP2 (light grey), NELF (rose taupe), DSIF (sand) and bound DNA transcription bubble (blue) are shown in surface representation. Location of sIBM and hIBM binding sites on INTS13-14 are indicated by orange circles. DNA path is extended by a thick blue line to visualize regions upstream or downstream of the transcription bubble, with a potential TF (orange) binding in the upstream region able to reach the TF contact sites on INT (dotted orange line). (L) Summary model of INT’s action on TF target genes. Red bars indicate potential points of regulation for INT recruitment via TFs (TF expression/DNA binding, TF IBM affinity for INT, INTS13 CMBM affinity for cleavage module).

We next inspected the published cryoEM map (EMDB 30473) of an INT preparation, which was purified from human cells directly^30^ for unexplained densities. We noticed a well resolved additional density section on a surface between INTS2-7 consistent with a short α-helix extended by an unstructured coil (so far modelled as unassigned poly-Ala chain). Since first identification of INT occurred after serendipitous co-immunoprecipitation with Flag-DSS1,^23^ we wondered whether this density could correspond to DSS1, which might have been inadvertently co-purified with INT. In support of this notion, DSS1 is 70aa short acidic protein that binds many complexes like the proteasome or TREX,^73^ and in these contexts forms a reminiscent α-helix with an unstructured linker.^74–76^ Indeed, we found that purified His_6_-tagged DSS1 efficiently and specifically binds purified INTS2-7 (Figure 7F). Furthermore, we could fit a large section of the protein sequence (aa35-66) into the unassigned density with high confidence (Figure 7G and Supplemental Figure 7M). Since it covers a hydrophobic patch on INTS7, DSS1 most likely has a stabilizing effect. Further, its binding to INT is probably mutually exclusive with its association to other complexes such as TREX, since the same DSS1 residues are involved in these interfaces.^74–76^ Thus, our data suggest that DSS1 is indeed a further stable INT subunit, which increases the total to 18 assigned components (INTS1-15, DSS1, PR65, PP2Ac).

### The TF binding sites of INT are accessible and located upstream of the transcription bubble on paused RNAP2

The published cryoEM density also features a large extended section of completely unmodeled density with low resolution that protrudes from INTS5 and stretches in a crescent shape behind the ICM (Figure 7H). In an attempt to define whether this density could represent ITFM and endogenous INTS15 that were co-purified to a lower extent with overexpressed INT from human cells, we superposed our completed INTS13-14-10-15-5 model onto INTS5, which is part of the published structure.^30^ Indeed, all additional subunits end up in the extra density and fit in remarkably well (Figure 7I). In support of this completed INT structure (Figure 7J), INTS10 is located close to the two CTD2s of INTS9-11 in agreement with crosslinks previously detected by MS.^13^ Furthermore, the C-terminus of INTS13’s α-helical domain, from which the CMBM protrudes, points towards the INTS11 surface where, according to our data, the INTS13-CMBM is located, and their distance (∼75 Å=min. 21 aa) could be easily bridged by the linker in INTS13 (85 aa). Finally, this placement proximal to the cleavage module would also be consistent with INTS10-13-14 contacting the RNA transcript upstream of the INTS11 cleavage site (distance to active site ∼115 Å=min. 30 nucleotides) as suggested by previous nucleic acid binding assays.^44,65^

Importantly, when we superpose this completed INT structure (Figure 7J) onto a structure of INT bound to PEC,^13^ the ITFM has its TF-binding surfaces freely accessible and is located close to the 5’ end of the coding DNA strand extruding from RNAP2 (Figure 7K). This means that the contact points for TFs on INTS13-14 in the context of PEC are arranged upstream of the transcription bubble, i.e. in a sterically favorable position that could promote simultaneous attachment of INT to promoter-bound TFs and promoter-proximally paused RNAP2 (Figure 7L). Taken altogether our data suggest a model, in which sequence-specific TFs in a regulated manner bind the ITFM directly at target promoters and thereby increase local concentrations of INT, which is favorably oriented to latch onto RNAP2 during pausing to prevent elongation.

## DISCUSSION

Transcription regulation via INT has been known to be required for inducible transcriptional programs for some time.^19,26,32,34,35,37,38^ Yet, it remained unclear how this coordinated response could be orchestrated by INT given that it binds promoter proximally paused RNAP2, which features on all RNAP2 transcribed genes.^2,3^ Despite efforts, no predictive common features were identified in INT-regulated coding genes (e.g. enriched TF motifs in promoters) or their transcripts (e.g. RNA sequence motifs or secondary structure elements).^14,17^ Even for the first and best established ncRNA targets of INT, UsnRNAs,^23^ the motif close to which INT-mediated 3’ end processing occurs (3’-box) is partially divergent, not found on all UsnRNA genes and not fully required for cleavage and termination.^44,77^ Indeed, recent studies suggest that INT co-localizes with paused RNAP2 indiscriminately on all active transcription units (TUs).^69,71^

In this study, we provide an explanation for the gene-specific and inducible regulation that has been observed for INT in the past. We define a diverse set of sequence-specific and developmentally important transcription factors that bind INT directly on two hubs of the INTS13-14 subcomplex. These TFs recruit INT to genomic loci, and by increasing local INT concentration on target genes, they co-regulate expression of these genes together with INT. The diverse nature of INT-bound TFs and their short linear (and evolutionary quite mobile) binding motifs have likely precluded their detection by conservation and enrichment analyses in the past.

We also demonstrate that these TF-INT connections are required for cellular stimulus response at the example of glucose starvation and primary cilia formation. Therefore, our data suggest that the molecular basis for the recently defined human ciliopathy that is caused by INTS13-CMBM mutations^37^ lies in the severed link between INT and TFs via loss of ITFM incorporation. Additionally, the direct INT binding that we defined for ZNF609, is likely also required for neurodevelopment in mammalian embryos based on the observation that both ZNF609 and INTS1/11 are necessary for cortical neuron migration in mice.^36^ Thus, the binding hubs that we identified are likely crucial for the function of INT during cell differentiation and embryo development.

Our analysis was focused on HEK293 cells, however it is probable that additional cell type- or state-specific transcription factors exploit the same binding sites on ITFM to recruit INT for targeted transcription regulation. For example, a previous study in a leukemia cell line suggested interaction of INTS13 with EGR1/2-NAB2 during differentiation.^35^ Although we could so far not confirm a direct interaction with INT (data not shown), it is possible that PTMs might be required for binding. Furthermore, additional, as of yet unidentified, contact sites for transcription effectors could exist on INT. For example, recent studies have suggested contacts to PRC2^39^, or a correlation of INT abundance on genes with histone mark H3K4me3.^78^ Importantly, we also found direct binding between INT and LEC, which constitutes the first UsnRNA-specific protein-protein contact of INT but the function of this interaction is still unclear.

For fast response to external cues, the recruitment of INT via TFs could be regulated at multiple levels (Figure 7L). Our data indicate that binding affinity of sIBMs could be increased by phosphorylation. Conversely, ITFM interaction with the core of INT could be reduced by phosphorylation. Furthermore, according to quantitative proteomic studies, the concentration of individual INTS differs substantially.^62,70^ In particular, INTS13-14 are two-fold more abundant than ICM subunits, in keeping with our and published ChIP data,^35^ where only half of all INTS13 ChIP peaks are matched by INTS11 peaks. In contrast, INTS10-15, which form the second anchor of ITFM to the INT core seem to be limiting, possibly explaining the crucial function of INTS13 CMBM for ITFM incorporation. Finally, TFs expression itself as well as accessibility of promoters for TF-binding at chromatin level could be adapted. In this context, it is relevant that we found the chromatin remodeler INO80 as a directly bound complex, which could signify that INT also exerts indirect regulation via modulation of the chromatin state.

How can we reconcile the presence of INT on all active TUs with our findings on gene-specific regulation? Several congruent studies have in the past shown that frequent premature transcription termination is a common feature of all coding genes.^6–8^ We therefore propose a scenario where INT binds PECs on all TUs transiently and in competition with elongation factors.^29,31^ INT could then terminate elongation-incompetent RNAP2 with higher likelihood in a quality control step, but still allow normal elongation to proceed, as suggested by recent publications.^20,21,71,80^ Strong TF-mediated recruitment of INT would in contrast lead to a more persistent attenuation of targeted genes by more efficient PEC termination. In this way, specific genes could remain poised for bursts of expression upon signaling cascades that remove the elongation block or put it in place.

## Supporting information

Supplemental Figures with Legends

## ACKNOWLEDGMENTS

We thank project students Valentin Erhard, Alexandra Burger, and İdil Itır Demiralp for support during early stages of the project, and Debbie van den Berg for generously sharing ZNF609 antibody. We acknowledge support by Functional Genomics Center Zurich through MS and next-generation sequencing (NGS), ScopeM ETH Zurich for confocal microscope access, and Swiss Light Source beamline scientists (PSI, Switzerland) for support during data collection. This work was supported by the SNSF through the NCCR “RNA & Disease” (to SJ, #141735, #182880, #205601), an ERC Consolidator grant funded via SERI (to SJ, #MB22.00064), an EMBO Young Investigators Award (to SJ, #4918), and a Ph.D. fellowship of the German Academic Scholarship Foundation (to KS).

## AUTHOR CONTRIBUTIONS

KS and SJ conceived the study, and obtained funding. AN, KS and SJ designed experiments, analyzed and interpreted data. KS supervised DD, performed biochemical experiments helped by DD, and proteomics and structural studies. AN performed NGS and IF experiments. Bioinformatic analyses of NGS data were supervised by JZ, and carried out by AN with support from CA and RM. SJ supervised the whole study. All authors contributed to manuscript writing.

## MATERIALS AND METHODS

### Antibodies

The following antibodies and dilutions were used for western blotting or immunofluorescence: horseradish peroxidase-linked anti-HA antibody (Roche, 1:5,000), rabbit anti-INTS4 (Bethyl Laboratories, A301-269A, 1:2,000), rabbit anti-INTS5 (Bethyl Laboratories, A301-268A, 1:5,000), rabbit anti-INTS8 (Bethyl Laboratories, A300-269A, 1:5,000), rabbit anti-INTS10 (Proteintech Group, 15271-1-AP, 1:2,000), rabbit anti-INTS11 (Bethyl Laboratories, A301-274A, 1:1,000), rabbit anti-INTS13 (Bethyl Laboratories, A303-575A, 1:5,000), rabbit anti-INTS14 (Bethyl Laboratories, A303-576A, 1:2,500), rabbit anti-C7ORF26 (Sigma-Aldrich, HPA052175, 1:1,000), rabbit anti-ZNF532 (Bethyl Laboratories, A305-442A, 1:1,000), rabbit anti-ZNF687 (Bethyl Laboratories, A303-279A, 1:1,000), rabbit anti-ZMYND8 (Proteintech, 11633-1-AP, 1:1,000), rabbit anti-ZEB1 (Proteintech, 21544-1-AP-150UL, 1:1,000), rabbit anti-HDGF (Proteintech, 11344-1-AP, 1:10,000), guinea pig anti-ZNF609 (1:1,000),^36^ mouse anti-INO80E (Santa cruz biotech, sc-515298, 1:1,000), rabbit anti-INO80 (Abcam, ab118787, 1:2,000), mouse anti-Strep-tag (IBA, 2-1507-001, 1:1,000) rabbit anti-CBP-tag (GenScript, A00635-40, 1:1,000), mouse anti-V5-tag (Proteintech Group, 66007-1-Ig, 1:5,000), mouse anti-β-actin (Proteintech Group, 60008-1-Ig, 1:20,000) horseradish peroxidase-linked anti-mouse IgG (Sigma-Aldrich, A9044, 1:5,000), horseradish peroxidase-linked anti-rabbit IgG (Sigma-Aldrich, A9169-2ML, 1:5,000), horseradish peroxidase-linked anti-guinea pig IgG (Sigma-Aldrich, A5545-1ML, 1:5,000), anti-ARL13B (for IF, Proteintech Group, 66739-1-Ig, 1:500), and anti-rabbit IgG (H+L) Alexa 488 (for IF, Invitrogen, A11008, 1:300).

### Plasmids

V5-SBP and λN-HA-tagged *Homo sapiens* (*hs*)INTS10, *hs*INTS13 and *hs*INTS14 full-length, truncations and mutant constructs as well as insect cell expression constructs for *hs*INTS4-*hs*INTS9-*hs*INTS11, *hs*INTS13-*hs*INTS14 and *hs*INTS10-*hs*INTS13-*hs*INTS14 have been described previously.^44^

cDNAs encoding full-length *hs*ZNF655, *hs*INO80E, *hs*HDGF, *hs*INO80, *hs*ICE2 *hs*ZMYND8, *hs*TSPYL2, *hs*DSS1, *hs*INTS3, *hs*INTS5, *hs*INTS6, *hs*INTS7, *hs*INTS11 and *hs*INTS12 were acquired from the human open reading frame library (hORFeome V5.1, ID: 10159, 11596, 6991, 53172, 9886, 55579, 10859, 11271, 2489, 5400 and 7700, hORFeome V8.1, ID: 101689522, 101686407 and 100002000, respectively). cDNAs of *hs*ZNF608, *hs*C7ORF26 and *hs*INTS2 were synthesized by reverse transcription from total HeLa cell RNA using gene-specific primers. cDNA encoding *hs*ZNF609 and *hs*ZEB1 were ordered from Genscript, while cDNAs encoding *hs*INTS1 and *hs*INTS8 were ordered as string DNAs from Thermo Fisher Scientific.

Mutations in coding sequences were introduced by the QuikChange method (Agilent) or round-the-horn mutagenesis using appropriate primers.^81^ All generated constructs were confirmed by sequencing.

To express V5-SBP-tagged or λN-HA-tagged *hs*ZNF655, *hs*INO80E, *hs*INO80, *hs*ZNF608, *hs*ZNF609, *hs*ZMYND8, *hs*HDGF, *hs*ICE2, *hs*TSPYL2, *hs*ZEB1, *hs*C7ORF26, *hs*INTS5, *hs*INTS8, *hs*INTS11 and *hs*INTS13 in human cells, the respective full length cDNA, truncations or mutants were inserted into the multiple cloning site of the pCIneo-V5-SBP or pCIneo-λN-HA plasmid to generate N-terminal fusion proteins or in the identical multiple cloning site in the pN3-V5-SBP or pN3-λN-HA to generate C-terminally tagged fusion constructs.^82,83^

For the expression of recombinant proteins in *E.coli*, cDNAs encoding full length or truncationed proteins were inserted into vectors of the pET-MCN series.^84^ *Hs*ZNF655, *hs*INO80E, *hs*HDGF, *hs*ZNF608, *hs*ZNF609, and *hs*INTS13 were inserted into the multiple cloning site of the pnEA-pM or pnEA-pG vector resulting in an N-terminal fusion with an MBP (pM) or GST (pG) tag cleavable by HRV 3C protease. Additionally, *hs*ZNF655 and *hs*DSS1 were inserted into the pnEA-CvH or pnYC-vG vector resulting in a C-terminal fusion protein with a TEV cleavable 6xHis tag (vH) or an N-terminal fusion with a GST tag cleavable by TEV protease (vG). cDNA encoding *hs*INTS13 (1-256Δ27-48) was inserted into the pnEA-pHB vector resulting in an N-terminal fusion with a hexahistidine-GB1 (pHB) tag cleavable by HRV 3C protease. To generate GFP fusion constructs the minimal *hs*INTS13-14 interacting sequences of *hs*ZNF655, *hs*ZNF608, *hs*ZNF609, *hs*HDGF, *hs*ZMYND8, *hs*TLE3, *hs*INO80E, *hs*ICE2, *hs*ZEB1, and *hs*ZEB2 were inserted in the multiple cloning site of a modified pnEA plasmid resulting in a fusion construct with an N-terminal GFP tag and a TEV protease cleavable C-terminal hexahistidine tag.

The coding sequence of *hs*INTS13ΔC (1-648), *hs*INTS10 or GFP was cloned into the pcDNA5/FRT/TO/n2SC or pcDNA5/FRT/TO/c2SC vector (Thermo Fisher Scientific) encoding for a tandem StrepII tag followed by a calmodulin binding peptide (CBP) at the N- or C-terminus.

Using primers with overhangs, full length *hs*INTS14 was C-terminally extended by a short SGS-linker followed by the INTS13-INTS14 interacting peptide of *hs*ZNF609 (INTS14-ZNF609 sIBM; aa 25-41).

For recombinant protein expression in insect cells full length or truncations of *hs*INTS1, *hs*INTS2, *hs*INTS3, *hs*INTS5, *hs*INTS6, *hs*INTS7, *hs*INTS8, *hs*INTS10 *hs*INTS12, *hs*INTS14-ZNF609 sIBM and *hs*C7ORF26 were cloned and assembled in multigene expression cassettes by ligation-independent cloning (LIC) using the MacroBac system.^85^ Genes of interest were PCR amplified introducing LIC-compatible overhangs. Modified pFastBac vectors that contain no tag or an N-terminal tandem StrepII (2S), MBP (M), GST (G) or decahistidine tag (10×His) that are cleavable by HRV 3C protease served as destination vectors. Vectors were linearized with *Ssp*I. Inserts and vectors were treated with T4 DNA polymerase (NEB), annealed and subsequently transformed into *E.coli* DH5α. For larger gene assemblies, expression cassettes were consecutively added by restriction digest followed by LIC assembly. Expression vectors containing 2S-*hs*INTS1–*hs*INTS12, 2S-*hs*INTS2–*hs*INTS7, 2S-*hs*INTS3–*hs*INTS6, 2S-*hs*INTS8–*hs*INTS5, M-*hs*INTS10(1-54)-2S-*hs*INTS5(1-234)-G-*hs*C7ORF26, 2S-*hs*INTS13-*hs*INTS14-ZNF609 sIBM and G-*hs*C7ORF26 were assembled. Bacmids were obtained after transformation of the expression vectors into *E.coli* DH10Bac cells and blue-white colony screening.^86^ Correct insertion of expression cassettes into bacmids was verified by PCR.

### Cell-lines

All human cells were cultured in Dulbecco’s modified Eagle’s medium (DMEM, Sigma-Aldrich) supplemented with 10% fetal calf serum (FCS), 100 U/mL penicillin, and 100 mg/mL streptomycin (Thermo Fisher Scientific). TAP cell lines were generated based on the HEK293 Flp-In T-REx system (Thermo Fisher Scientific). HEK293 cell lines with tetracycline inducible TAP-tagged baits were generated by co-transfection of 0.1 µg of plasmid containing the TAP-tagged bait (*hs*INTS13ΔC; *hs*INTS10; GFP) in a pcDNA5/FRT/TO/n2SC or pcDNA5/FRT/TO/c2SC vector with 0.9 µg of pOG44 plasmid into HEK 293 Flp-In T-REx cells (Invitrogen). Cells were selected for 2 weeks in DMEM containing 100 µg/mL hygromycin and 15 µg/mL blasticidin. Expression of the desired constructs upon tetracycline induction was verified by Western blotting.

### Tandem affinity purification

6 × 10^6^ cells were seeded per 15 cm plate in DMEM. After 24 h, medium was exchanged to DMEM supplemented with tetracycline. To achieve comparable expression levels with the endogenous protein and between the target protein and the GFP control the following concentrations were used: INTS13-TAP: 0.05 µg/mL for *hs*INTS13ΔC and 0.02 µg/mL for GFP cell lines; INTS10-TAP: 0.02 µg/mL for *hs*INTS10 and 0.013 µg/mL for GFP cell lines. 48 h after induction cells were harvested in PBS containing 0.5 mM EDTA and pelleted at 900 *g* for 5 min at 4 °C. In case of glucose starvation, the medium was changed to DMEM without glucose but containing tetracycline 24 h after induction. For heat shock the plates were transferred for 2 h before harvesting to an incubator at 42 °C. For EGF treatment, cells were induced with tetracycline for 48 h in DMEM with only 0.5% FCS before addition of 100 ng/mL EGF 1 h prior to harvesting.

Cells were lysed in 20 mM HEPES pH 7.6, 150 mM NaCl, 0.5% NP-40, 2 mM DTT supplemented with 1×cOmplete EDTA-free protease inhibitor cocktail (Roche), 1×PhosSTOP phosphatase inhibitor (Roche), 5 µg/mL DNase I (AppliChem) and 200 µg/mL RNase A (Qiagen) using 4 cycles of 15 s sonification at 30% duty cycle and 30% output control. The lysate was cleared by centrifugation at 10,000 *g* for 30 min at 4 °C.

The lysate was incubated with 100 µL pre-equilibrated StrepTactin sepharose beads (IBA) for 1 h at 4 °C while rotating on a wheel. Beads were washed twice with buffer A (20 mM HEPES pH 7.6, 150 mM NaCl, 0.1% NP-40, 5 mM EGTA, 2 mM DTT) and twice with buffer B (20 mM HEPES pH 7.6, 150 mM NaCl, 0.1% NP-40, 2 mM DTT). Bound complexes were eluted with four times 100 µL of buffer C (20 mM HEPES pH 7.6, 150 mM NaCl, 0.1% NP-40, 2 mM CaCl_2_, 5 mM biotin, 2 mM DTT). The eluates were pooled and incubated with 50 µL pre-equilibrated calmodulin sepharose 4B (GE Healthcare) for 1 h at 4 °C while rotating on a wheel. Beads were washed three times with buffer D (20 mM HEPES pH 7.6, 150 mM NaCl, 0.1% NP-40, 2 mM CaCl_2_, 2 mM DTT). Proteins were eluted with three times 50 µL of buffer A with 1 M NaCl and pooled for analysis by SDS-PAGE followed by silver staining or mass-spectrometry.

### LC-MS/MS sample preparation and data acquisition

Proteins were precipitated with trichloroacetic acid (TCA; Sigma-Aldrich) at a final concentration of 5% and washed twice with ice-cold acetone. Protein pellets were resuspended in 45 µL digestion buffer (triethylammonium bicarbonate (TEAB), pH 8.2), reduced and alkylated. 500 ng of sequencing grade trypsin (Promega) were added for digestion carried out overnight at 37 °C. The samples were dried to completeness and re-solubilized in 20 µL of MS sample buffer (3% acetonitrile, 0.1% formic acid).

LC-MS/MS analysis for *hs*INTS13ΔC TAP sample under normal growth conditions or stress conditions was performed on an Orbitrap Fusion Lumos (Thermo Scientific) equipped with a Digital PicoView source (New Objective). For *hs*INTS10, LC-MS/MS analysis was performed on an Orbitrap Exploris 480 mass spectrometer (Thermo Fisher Scientific) equipped with a Nanospray Flex ion source (Thermo Fisher Scientific). All machines were coupled to an M-Class UPLC (Waters). Solvent composition of the two channels was 0.1% formic acid for channel A and 99.9% acetonitrile in 0.1% formic acid for channel B. Column temperature was 50 °C. For each sample 3 µL (*hs*INTS13ΔC under normal growth conditions) or 4 µL (*hs*INTS13ΔC under stress conditions) of peptides were loaded on a commercial ACQUITY UPLC M-Class Symmetry C18 Trap Column (100 Å, 5 µm, 180 µm x 20 mm, Waters) connected to a ACQUITY UPLC M-Class HSS T3 Column (100 Å, 1.8 µm, 75 µm x 250 mm, Waters). The peptides were eluted at a flow rate of 300 nL/min. After a 3 min initial hold at 5% solvent B, a gradient from 5 to 22% solvent B over 80 min and 22 to 32% B over additional 10 min was applied. For the INTS10 TAP sample 4 µL were loaded on a commercial nanoEase MZ Symmetry C18 Trap column (100 Å, 5 µm, 180 µm x 20 mm, Waters) followed by a nanoEase MZ C18 HSS T3 column (100 Å, 1.8 µm, 75 µm x 250 mm, Waters). The peptides were eluted at a flow rate of 300 nL/min. After a 3 min initial hold at 5% solvent B, a gradient from 5 to 35% solvent B over 90 min. The column was cleaned after the run by increasing to 95% solvent B for 10 min prior to re-establishing loading conditions. Samples were measured in randomized order. The mass spectrometer was operated in data-dependent mode (DDA) with a maximum cycle time of 3 s, with spray voltage set to 2.5 kV, funnel RF level at 40%, heated capillary temperature at 275 °C, and Advanced Peak Determination (APD) on. Full-scan MS spectra (300−2,000 m/z for the *hs*INTS13ΔC sample under normal growth conditions, 300−1,500 m/z for the *hs*INTS13ΔC samples under stress conditions or 350−1,200 m/z for the *hs*INTS10 sample) were acquired at a resolution of 120,000 at 200 m/z after accumulation to an automated gain control (AGC) target value of 500,000 for a maximum injection time of 50 ms for the *hs*INTS13ΔC samples or accumulation to a target value of 3,000,000 or for a maximum injection time of 45 ms. Precursors with an intensity above 5,000 were selected for MS/MS. Ions were isolated using a quadrupole mass filter with 1.6 m/z for the *hs*INTS13ΔC sample under normal growth conditions, for 0.8 m/z for the *hs*INTS13ΔC samples under stress conditions and 1.2 m/z for the *hs*INTS10 sample isolation window and fragmented by higher-energy collisional dissociation (HCD) using a normalized collision energy of 35% for *hs*INTS13ΔC or 30% for *hs*INTS10. For *hs*INTS13ΔC samples fragments were detected in the linear ion trap with the scan rate set to rapid, the automatic gain control set to 10,000 ions, and the maximum injection time set to 50 ms. Charge state screening was enabled, and singly, unassigned charge states as well as charge states higher than seven were excluded. For *hs*INTS10, HCD spectra were acquired at a resolution of 15,000 with maximum injection time set to “Auto” and AGC set to 100,000 ions. Charge state screening was enabled such that singly, unassigned and charge states higher than six were rejected. Precursor masses previously selected for MS/MS measurement were excluded from further selection for 25 s (for *hs*INTS13ΔC under normal growth conditions) or 20 s (for *hs*INTS13ΔC under stress conditions and *hs*INTS10), applying a mass tolerance of 10 ppm. Data was acquired using internal lock mass calibration on m/z 371.1012 and 445.1200.

### Analysis of LC-MS/MS data

Peptide identification and quantification was performed with MaxQuant (version 2.0.2.0) using Andromeda as search engine.^87^ The human proteome subset of UniProt (UP000005640) combined with the contaminant database from MaxQuant was searched and the protein and peptide FDR values were set to 0.01.

Statistical analysis was done in Perseus (version 1.6.15.0).^88^ Results were filtered to remove reverse hits, contaminants and peptides found in only one sample. Missing values were imputed from normal distribution. Label free quantification (LFQ) results were exported from Perseus and visualized using statistical computing language R. Venn diagrams were generated using InteractiVenn.^89^

### Recombinant protein expression in *E.coli*

The following constructs were expressed separately or together in *E.coli* BL21 Star (DE3) cells (Invitrogen): MBP/6xHis-GB1-*hs*INTS13 (VWA: 1-256, β-barrel: 257-389, α-helical: 413-564, VWAΔloop: 1-256Δ27-48), MBP/GST-*hs*ZNF655 (1-125, 1-56, 57-125, 93-119), MBP-*hs*ZNF608 (1-51, 1-140, 24-51, 24-140), MBP-*hs*ZNF609 (25-41, 25-41_I35A, 25-41_I36A), GST-*hs*INO80E (1-244, 1-61, 62-211, 212-244), 6xHis/6xHis-GST-*hs*HDGF (1-145, 125-145, 117-145) and *hs*DSS1-6xHis. Numbers in brackets specify domain boundaries. Cells were grown at 37 °C in LB medium with appropriate antibiotics until OD_600_=0.5 was reached, then 1 mM isopropyl-β-D-thiogalactopyranoside (IPTG) was added and cells were incubated overnight at 20 °C.

GFP-6xHis(C)-tagged *hs*ZNF655 (93-119), *hs*ZNF608 (24-40), *hs*HDGF (130-146), *hs*ZMYND8 (702-720), *hs*ZNF609 (25-41), *hs*HDGF S2E (130-146), *hs*TLE3 (238-255), *hs*INO80E (229-244), *hs*ICE2 (568-585), *hs*ZEB2 (33-50), *hs*ZEB1 (28-45) were expressed in *E.coli* Rosetta (Sigma Aldrich). Cells were grown at 37 °C in LB medium with appropriate antibiotics until OD600 = 0.5 was reached, then 1 mM isopropyl-β -D-thiogalactopyranoside (IPTG) was added and cells were incubated for 3 hours at 37 °C.

### Recombinant protein expression in insect cells

Protein complexes (2S-*hs*INTS1–*hs*INTS12, 2S-*hs*INTS2–*hs*INTS7, 2S-*hs*INTS3–*hs*INTS6, 2S-*hs*INTS8–*hs*INTS5, G-*hs*INTS15, 2S-*hs*INTS13-*hs*INTS14, 2S-*hs*INTS13-*hs*INTS14-ZNF609 sIBM, *hs*INTS10-(2S)-*hs*INTS13-*hs*INTS14, M-*hs*INTS10 1-54-2S-*hs*INTS5 1-234-G-*hs*INTS15 or 10×His-*hs*INTS4-*hs*INTS9-(2S)-*hs*INTS11) for structure determination and interaction studies were expressed in insect cells. Bacmid DNA isolated from *E.coli* DH10Bac cells was used to transfect Sf9 cells (Thermo Fisher Scientific) using EscortIV transfection reagent (Merck) growing in SF-4 Baculo Express ICM medium (BioConcept) to generate baculovirus. After 2 rounds of virus amplification in Sf9 cells, protein expression was carried out in HighFive cells (Thermo Fisher Scientific) at 130 rpm and 27 °C for 48 h. Cells were harvested by centrifugation at 900 *g* for 10 min at 4 °C.

### Protein purification

To purify *hs*INTS13 1-256Δ27-48 in complex with *hs*ZNF655 93-119, the cell pellet was resuspended in lysis buffer 1 (50 mM HEPES pH 7.5, 200 mM NaCl, 20 mM imidazole, 2 mM β-mercaptoethanol) supplemented with 1×cOmplete EDTA-free protease inhibitor cocktail, 1 mg/mL lysozyme and 5 µg/mL DNase I and lysed using a microfluidizer (Microfluidics). The cleared lysate was filtered (0.45 µm) and bound to glutathione sepharose 4B resin (Amersham). Beads were washed with lysis buffer 1 and bound proteins were eluted in lysis buffer 1 supplemented with 25 mM glutathione. The eluate was incubated overnight with TEV-protease (homemade). The protein complex was further purified over a Ni-charged chelating column (GE Healthcare) and eluted by a linear gradient to 500 mM imidazole. The 6xHis-GB1 tag was removed by overnight cleavage using HRV 3C protease (homemade) during dialysis against lysis buffer. The cleaved tags were removed by re-binding to a Ni-charged chelating column. Finally, size-exclusion chromatography was performed using a Superdex75 column (GE Healthcare) pre-equilibrated in gel filtration buffer (10 mM HEPES at pH 7.5, 200 mM NaCl, 2 mM DTT). The complex was immediately used to set up crystallization plates.

G-*hs*INTS15 and 2S-*hs*INTS13-*hs*INTS14-ZNF609 sIBM were expressed in insect cells. Cell pellets were resuspended in lysis buffer 2 (50 mM HEPES pH 7.5, 200 mM NaCl, 5% glycerol, 2 mM DTT) supplemented with 1×cOmplete EDTA-free protease inhibitor cocktail and 5 µg/mL DNase I and lysed by sonication or by using a microfluidizer. Cell lysates were cleared and subsequently filtered (0.45 µm).

G-*hs*INTS15 was applied to pre-equilibrated glutathione sepharose 4B resin. Beads were washed with lysis buffer 2 and bound proteins were eluted in lysis buffer 2 supplemented with 25 mM glutathione (Carl Roth). The sample was applied to a PD-10 desalting column (Cytiva) and eluted in lysis buffer 2 to remove glutathione from the sample. Subsequently, the sample was used for biochemical assays or flash-frozen in liquid nitrogen and stored at −80 °C.

2S-*hs*INTS13-*hs*INTS14-ZNF609 sIBM was applied to a pre-equilibrated StrepTrap column (GE Healthcare), washed with lysis buffer 2, and protein complex was eluted in lysis buffer 2 containing 2.5 mM D-desthiobiotin (Merck). Protein tags were removed by overnight cleavage with HRV 3C protease. The proteins were subsequently purified over a heparin column (GE Healthcare). Complexes were eluted by a linear gradient to 1 M NaCl. Finally, the complexes were purified by gel filtration (Superose 6, GE Healthcare) pre-equilibrated in gel filtration buffer. The proteins were either used to set up crystallization plates, in biochemical assays or flash-frozen in liquid nitrogen and stored at −80 °C.

To purify GFP-tagged INTS13-INTS14 interacting peptides and *hs*DSS1, the cell pellets were resuspended in lysis buffer 1 supplemented with 1×cOmpleate EDTA-free protease inhibitor cocktail, 1 mg/mL lysozyme and 5 µg/mL DNase I and lysed using a microfluidizer. The cleared lysate was filtered (0.45 µm) and bound to a Ni-charged chelating column and eluted by a step elution to 500 mM imidazole. For the GFP-tagged samples, the eluates were incubated overnight with TEV-protease and dialyzed against lysis buffer 1. Cleaved tags were removed by re-binding to the Ni-charged chelating column. The flow throughs were collected, concentrated and further purified using a Superdex 75 column pre-equilibrated in gel filtration buffer. For *hs*DSS1, the eluate from the Ni-column was diluted to 100 mM NaCl and further purified using a Q-column (GE Healthcare). Bound proteins were eluted by a linear gradient to 1 M NaCl. Fractions containing *hs*DSS1 were pooled, concentrated and flash-frozen in liquid nitrogen for storage at −80 °C.

Insect cells expressing recombinant 2S-*hs*INTS2-*hs*INTS7 or 2S-*hs*INTS8-*hs*INTS5, were lysed using a microfluidizer in lysis buffer 2 supplemented with 1×cOmplete EDTA-free protease inhibitor cocktail and 5 µg/mL DNase I. The lysate was cleared and filtered (0.45 µm). Complex was purified using a StrepTrap column (GE Healthcare), washed with lysis buffer 2, and protein complex was eluted in lysis buffer 2 containing 2.5 mM D-desthiobiotin. In case of 2S-*hs*INTS8-*hs*INTS5, the complex was further purified using a heparin column. The sample was diluted to 100 mM NaCl, bound to the column, washed and proteins were eluted by a linear gradient to 1 M NaCl. Protein complexes were pooled, concentrated and flash-frozen in liquid nitrogen.

2S-*hs*INTS13-*hs*INTS14 and *hs*INTS10-(2S)-*hs*INTS13-*hs*INTS14 were purified as previously described.^44^

### Crystallization and structure determination

Initial screens were carried out using the sitting drop vapor diffusion method using 7.8 mg/mL of the *hs*INTS13(1-256Δ27-48)-*hs*ZNF655(93-119) complex or 10.3 mg/mL of the *hs*INTS13-*hs*INTS14-ZNF609 sIBM complex. The complex (200 nL) was added to 200 nL of reservoir solution. For the *hs*INTS13(1-256Δ27-48)-*hs*ZNF655(93-119) complex crystals appeared after several days and the best diffracting crystal was obtained in 100 mM MES pH 6.0, 100 mM MgCl_2_ and 8% (w/v) polyethylene glycol 6,000. Crystals were cryoprotected using reservoir solution supplemented with 15% (v/v) glycerol and flash frozen in liquid nitrogen.

For the *hs*INTS13-*hs*INTS14-ZNF609 sIBM complex crystals appeared within one day in many different conditions. The best crystals grew in 100 mM HEPES pH 6.5, 0.9 M sodium malonate and 0.25% (v/v) Jeffamine ED-2003. Crystals were cryoprotected using reservoir solution supplemented with 15% (v/v) glycerol and flash-frozen in liquid nitrogen.

Diffraction data were recorded on a PILATUS 6M detector at the PXIII beamline of the Swiss Light Source (SLS) at a temperature of 100 K. Data were processed using XDS (version February 5, 2021).^90^ Initial phase information was achieved by molecular replacement in Phenix (version 1.15.2-3472)^91^ using only the VWA domain of *hs*INTS13 or the apo *hs*INTS13-*hs*INTS14 structure as search model, respectively (PDB ID: 6SN1).^44^ The resulting map was of sufficient quality to build the structure. Iterative cycles of model building in COOT (version 0.8.9.2)^92^ and refinement performed with Phenix against the high-resolution native dataset were then used to finalize the structure.

The final model contains 3 copies of the *hs*INTS13(1-256Δ27-48)-*hs*ZNF655(93-119) complex in the asymmetric unit cell. All copies are overall very similar. The most complete copy of the complex contains residues −2-256 of *hs*INTS13 and residues 94-115 of *hs*ZNF655, except for a few disordered surface loops that were omitted from the model (INTS13 residues 22-29 and 193-195). Overall, the model of INTS13 is very similar to the INTS13-INTS14 structure.

The final model of the *hs*INTS13-*hs*INTS14-ZNF609 sIBM complex contains residues −1–564 of INTS13, 2–512 of INTS14 and 28–40 of ZNF609, except for a few disordered surface loops that were omitted from the model (INTS13 residues 34–40, 294–311, 517–522, and INTS14 residues 120-139, 286–295 as well as the C-terminal linker to the ZNF609 sIBM). Overall, the model is very similar to the apo structure, apart from two loops. We remodeled a short loop on top of the VWA domain of INTS14 (residues 116-139) and a loop of INTS13 close to the ZNF609 peptide that becomes ordered upon binding (residues 268–278).

### Co-affinity purification assays

For co-APs from *E.coli* or insect cells, pellets were generated as described above. Cell pellets were resuspended in lysis buffer 2 supplemented with 1×cOmplete EDTA-free protease inhibitor cocktail, 1 mg/mL lysozyme (*E.coli* pellets), and 5 µg/mL DNase I and lysed by sonication (*E.coli* pellets: 60 s sonification at 80% duty cycle and 30% output control, insect cell pellets: 60 s sonification at 30% duty cycle and 30% output control). The cleared lysates or purified components were mixed with 50 µL of glutathione sepharose 4B resin, StrepTactin sepharose or Ni-NTA agarose (Qiagen). 2S-GST, Strep-MBP or 6xHis-MBP served as negative controls. Proteins were incubated for 60 min on a rotating wheel at 4 °C, beads were washed four times with 700 µL lysis buffer 2. To test if presence of RNA can compete with TF binding, the beads were washed 3 more times with 100 µL lysis buffer 2 supplemented with 100 µg/mL polyU RNA (Merck). The bound complexes were eluted in lysis buffer 2 containing 25 mM glutathione or 5 mM biotin (Merck), respectively. For analysis 2×protein sample buffer was added to the eluates, proteins were separated by SDS–PAGE and detected by Coomassie staining or Western blotting.

For co-APs from human cells, 2.7 × 10^6^ HEK293T cells were seeded in 10 cm plates in 10 mL DMEM and transfected 24 h later using the calcium phosphate method. To express V5-SBP and HA-tagged proteins, cells were transfected with 20 µg of total plasmid and medium was changed to fresh DMEM the following day. V5-SBP-MBP or V5-SBP served as negative controls. Two days after transfection, cells were lysed for 10 min on ice in NET buffer (50 mM Tris-HCl pH 7.5, 150 mM NaCl, 0.1% Triton X-100, 10% glycerol, 2 mM DTT) supplemented with 1×cOmplete EDTA-free protease inhibitor cocktail, 5 µg/mL DNase I and 200 µg/mL RNase A (Qiagen). Followed by a mild sonication (10 strokes, 20% duty cycle, output control 2), cell lysates were centrifuged at 16,000 *g* for 15 min at 4 °C. The cleared lysate was rotated for 1 h at 4 °C in presence of 50 µL of streptavidin sepharose (50% slurry, GE Healthcare). Beads were washed three times with NET buffer. Bound proteins were eluted with 100 µL 2×protein sample buffer. For further analysis, proteins were separated by SDS-PAGE and detected by Western blotting.

### Fluorescence polarization assays

Binding reactions were carried out with 100 nM of the respective GFP-tagged peptide in binding buffer (50 mM HEPES pH 7.5, 200 mM NaCl, 2 mM DTT). INTS13-14 complex at concentrations ranging from 0.25 nM to 4 μM were briefly incubated with the GFP-tagged peptide in a black 384-well plate (Greiner) in a total reaction volume of 30 µL. FP was determined with a CLARIOstar plus microplate reader (BMG Labtech, Software version 6.10) by excitation at 482 nm and detection at 530 nm. Measurements on the same sample were repeated up to five times and all samples were prepared in triplicates. After baseline substraction, FP values were normalized to 1 using Microsoft Excel. Mean values of experimental triplicates and their standard deviation were plotted against the protein concentration and fitted using GraphPad Prism (Version 9.4.1) to a Hill equation:^93^

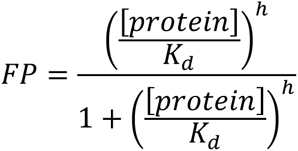

### Structure prediction using AlphaFold2

Structure predictions of the following complexes were carried out: *hs*INTS10-*hs*INTS14(1-210), *hs*INTS5(27-253)*-hs*INTS10(1-54)-*hs*INTS15, *hs*INTS9-*hs*INTS11-*hs*INTS13(649-700). Full length sequences or numbers in brackets specifying the boundaries were used as inputs. Models were predicted using AlphaFold2 implemented in ColabFold.^94,95^ Default settings were used and no additional constraints were applied.

### Model building and re-analysis of published cryoEM density maps

To build the INTS13-CMBM, the AF2 model for INTS9-INTS11-INTS13(aa649-690) was placed in the cryoEM density (EMDB 30473-2)^30^ and manually adjusted to fit the density using Coot. For DSS1, the model was built manually into cryoEM density (EMDB 30473-3)^30^ using Coot. To generate a model of INT containing INTS10-13-14-15, an AF2 model of full length INTS5 was superposed on PDB 7CUN^30^ to place the missing N-terminus of INTS5. Afterwards, the INTS5(aa27-253)-INTS15-INTS10(aa1-54) model was superposed on the INTS5 N-terminus. Subsequently, the INTS10-14(aa1-210) model was superposed on INTS10 and finally the INTS13-14 crystal structure (PDB 6SN1)^44^ was superposed on the VWA domain of INTS14.

### Sequence searches and alignments

Sequences of target proteins from *Homo sapiens* and orthologs were retrieved from Uniprot (http://www.uniprot.org) and aligned using the MUSCLE webserver^96^ from within JALVIEW.^97^ Positional conservation and similarity scores were calculated using the ESPRIPT webserver^98^ with default settings and converted to alignment figures manually.

### RNA-sequencing library preparation and analysis

All RNAseq experiments were carried out in biological duplicates. Per well of a 6-well plate, 75,000 HEK293T cells were seeded in 2 mL DMEM. The following day, growth medium was exchanged and gene-specific siPOOLs or a control siPOOL were transfected using 4 µL RNAiMax to a final concentration of 5 nM. After 24 h, transfections were repeated and cells were left to proliferate for another 48 hours. For starvation conditions, growth medium was exchanged to DMEM without glucose (Gibco) 24 h after the second transfection and cells incubated for another 24 h. For harvesting, medium was aspirated and cells washed with PBS before addition of trizol (Ambion) and RNA was extracted using phenol/chloroform. After DNase I (Qiagen) treatment, ribosomal RNAs were depleted and sequencing libraries were prepared using the Illumina TruSeq Stranded Total RNA kit. Libraries were sequenced on an Illumina NovaSeq 6000 using 100 bp single-end reads at a depth of 40×10^6^ reads per library. For proteins of which no specific antibodies are available (ZNF655 and ZNF608), validation of siPOOL depletions was achieved by reverse transcription followed by RT-qPCR. In brief, following DNase I treatment, 500 ng of RNA were reverse transcribed using random hexamer primers (Thermo Fisher Scientific) and AffinityScript reverse transcriptase (Agilent). cDNA products were diluted in water (1:5) and 5 µl of diluted cDNA was used for each qPCR reaction carried out using KAPA SYBR FAST master mix (Roche). Relative expression levels for each gene were determined based on the 2−ΔΔCT method normalizing against 7SK RNA.

For RNAseq data analysis, we used the workflow manager Snakemake^99^ to generate an analysis pipeline that starts with raw fastq files, performs initial quality control with FastQC (v0.11.5) followed by adapter trimming with Cutadapt (v1.15),^100^ using a mimimum read length of 30 bp. STAR-based (v2.4.1)^101^ alignment was performed to hg38 using the Gencode v34^102^ annotation, based on the following parameters: --outFilterMulitmapNmax 20 --quantMode GeneCounts --outFilterMismatchNmax 999 --outFilterMismatchNoverReadLmax 0.04 --alignIntronMin 20 --alignIntronMax 1000000 --alignMatesGapMax 1000000 --alignSJoverhangMin 8 --alignSJDBoverhangMin 1. Differential gene expression analysis was performed using DESeq2 (v1.40.1), following the workflow as described in the vignette using the design formula “∼ condition”.^103^ Genes were filtered out if they had less than 5 reads on average in either one of the conditions after read count normalization. Genes were considered misregulated if they had an absolute log_2_ fold change (L2FC) of at least 0.58 and a p-adjusted value of less than 0.05 using FDR correction for n=2 biological replicates and >17,000 observations (genes after filtering) per analysis. Heatmaps were produced using the Pheatmap R Package (v1.0.12). The GeneOverlap R package (v1.34.0) was used to assess and visualize gene overlaps and perform statistical analyses (odds ratio, Fisher’s-exact test). Gene ontology analysis was performed using the R package for Enrichr (v3.2) GO-biological pathways^104^ and bar graphs of the significant GO-terms were visualized using GraphPad Prism. GO terms were considered significantly enriched if they had a p-value of <0.05. Comparisons of normalized read counts and L2FCs and their statistical analysis were also performed in GraphPad Prism.

Statistical significance between normalized read counts and L2FCs of genes of the same class (e.g. all starvation response genes starved *vs.* normal conditions) was calculated using paired ratio-based t-tests. Comparison of different genes sets (e.g. INTS13-dependent *vs.* INTS13-independent) was based on unpaired t-tests.

### Chromatin immunoprecipitation library preparation and analysis

Antibodies used for ChIPseq were validated by IF in HEK293T upon control or target protein siRNA depletion. For each condition, 20×10^6^ HEK293T cells were cultured on a 10 cm dish in 10 mL DMEM. Cells were fixed using 1% formaldehyde by adding 10 mL of DMEM (no serum or antibiotics) to the cells at room temperature and shaking them gently for 5 min. The crosslinking reaction was quenched with 125 mM glycine and cells were washed with ice cold PBS. Cell lysis was achieved by adding 5 mL swelling buffer (5 mM PIPES pH 8, 85 mM KCl, 0.5% IGEPAL, cOmplete EDTA free protease inhibitor cocktail), after which cells were scraped off the dish and transferred to a falcon tube, where they were incubated on ice for 15 min. At this point, the lysed cells were pooled into one tube. The nuclei were pelleted by centrifugation at 2000 *g* for 5 min at 4 °C and the nuclei pellet was resuspended in 600 µL ice cold RIPA-1%SDS buffer (PBS, 1% IGEPAL CA-630, 0.5% sodium deoxycholate, 1% SDS, cOmplete EDTA free protease inhibitor cocktail). Chromatin was sonicated on a Diagenode Bioruptor300 for 30 min (30 s ON, 30 s OFF) on the high setting. After sonication the chromatin was cleared by centrifugation at max speed for 15 min. 10% of the resulting chromatin was taken as an input sample and the remaining chromatin was divided equally for each ChIP experiment. The chromatin was then diluted 10 times with RIPA-0% SDS (PBS, 1% IGEPAL CA-630, 0.5% sodium deoxycholate, cOmplete EDTA free protease inhibitor cocktail) such that the ChIP can be performed in classical RIPA buffer. Each sample was pre-cleared using 10 µL of protein A dynabeads (Thermo Fisher Scientific), before 10 µL of antibody (anti-INTS13, anti-INTS11, anti-ZEB1) were added and incubated overnight on a rotating wheel at 4 °C. For ZNF609 ChIP, 30 µL of antibody were used.^36^ The following day 20 µl protein A Dynabeads were used to recover the protein/DNA complexed by incubation on a rotating wheel for 4 h at 4 °C. Complexes were then washed at room temperature with the following buffers: Low-salt buffer (16.7 mM Tris-HCl pH 8, 0.167 M NaCl, 0.1% SDS, 1% Triton X-100, two washes), high-salt buffer (16.7 mM Tris-HCl pH 8, 0.5 M NaCl, 0.1% SDS, 1% Triton X-100, one wash), LiCl buffer (0.25 M LiCl, 0.5% Na-deoxycholate, 1 mM EDTA pH 8, 10 mM Tris-HCl pH 8, 0.5% IGEPAL, two washes), TE buffer (10 mM Tris-HCl pH 8, 5 mM EDTA pH 8, two washes). Protein/DNA complexes were eluted off the beads using 30 µL elution buffer (1% SDS, 100 mM NaHCO_3_) by incubation at 37 °C and 900 rpm shaking for 30 min. Inputs were also incubated with elution buffer alongside the ChIP samples. The ChIP samples were then placed on a magnetic rack and the supernatant was transferred to a fresh tube. In order to reverse the crosslinks, all samples were incubated with proteinase K mix (15 µL 1 M Tris-HCl pH 8, 15 µL 5 M NaCl, 7.5 µL 0.5 M EDTA pH 8, 1.2 µL proteinase K (20 mg/mL, AppliChem)) at 50 °C and 1100 rpm shaking for 3 h then 65 °C overnight. The next morning, each sample was treated with 10 µg of RNase A for 1 h at 37 °C. DNA was extracted using phenol/chloroform and quantified using the Qubit dsDNA Broad Range assay (Thermo Fisher Scientific). At least 20 µg of DNA were used for library preparation using the NEBNext Ultra DNA library prep kit. Libraries were sequences on an Illumina NovaSeq 6000 with 150 bp paired-end reads.

For ChIP-seq data processing and analysis, we used a modified version of the ATAC-seq Snakemake pipeline described in^105^, which excludes ATAC-seq specific steps. Briefly, we first perform FastQC quality control, then Trimmomatic (v0.36) adapter trimming using the parameters ILLUMINACLIP:NexteraPE-PE.fa:1:30:4:1:true TRAILING:3 MINLEN:20^106^, followed by Bowtie2 (v2.3.0) alignment to hg38 with -X2000 (maximal fragment length) and -very-sensitive as parameters^107^. After alignment, various cleaning steps were performed including removal of mitochondrial reads, reads with a mapping quality below 10 and duplicate reads with Picard tools (v2.9.0). MACS2 (v.2.1.1)^108^ was then used for peak calling with a minimum q-value of 0.1, followed by ChIPseeker for peak annotation (v1.2.0)^109^. bamCompare of the deeptools suite (v3.5.0)^110^ was used to normalize ChIP bigwigs against input sample bigwigs. The deeptools suite was used to further analyze and visualize all ChIP-seq data. computeMatrix was used to calculate ChIP coverage scores across the genome from normalized bigwig files, relative to transcription start sites. plotHeatmap was used to generate heatmaps from matrices and plotProfile was used to plot the ChIP profile of various samples at different gene sets. All tools were used in their default modes. Bedfiles of different gene sets were obtained from the UCSC table browser.^111^ To define the promoters of genes, we annotated the ChIPseq peaks of INTS13 and INTS11using ChIPseeker (parameters: up to 5kb upstream of the TSS).

### Immunofluorescence

HEK293T cells were seeded at 250,000 cells per well in 2 mL DMEM in 6-well plates on glass cover slips. The following day, cells were transfected with gene-specific siRNAs or a control siRNA (MISSION siRNA Universal Negative Control #1, Merck) to a final concentration of 45 nM and 400 ng of the respective rescue plasmid with 12 µL of Liopfectamine 2000. The transfections were repeated once more 24 h later. One day after the second transfection, growth medium was exchanged to a glucose-free DMEM and incubated for 24 h. For harvest, cover slips were transferred to PBS for 5 min and the remaining cells were harvested in protein sample buffer for validation of knockdown and overexpression by Western blot. After washing, coverslips were fixed in ice cold methanol at −20 °C for 5 min. All following steps after fixation were performed at room temperature. Cells were washed twice in PBS for 5 min each, then once for 10 min in BSA/PBS (2% BSA in PBS). After blocking in blocking solution (10% goat serum in BSA/PBS), cells were incubated with primary antibody, which was diluted in blocking solution for 1 h. Coverslips were then washed three times with BSA/PBS for 4 min each, before incubation with secondary antibody diluted in blocking solution for 1 h. Next, coverslips were washed three times in PBS for 4 min each, after which they were briefly incubated in IF detergent (0.1% TX-100, 0.02% SDS in PBS) before post-fixation in 4% paraformaldehyde (in PBS pH 7.4) for 20 min. After a 5 min wash in PBS, cells were stained with DAPI (1:2000 of 1 mg/mL in PBS), washed once in PBS for 5 min and then mounted on glass slides with 1.5 µL vectashield (ReactoLab).

Images were collected on a Confocal Zeiss LSM 780 microscope using the Zeiss ZEN software. Three Z-stacks were collected per condition such that the entire cell layer was imaged from top to bottom, with each slice set to 0.7 µm in height. Nuclei and cilia were counted manually by scrolling through Z-stacks and using the Cell Counter plugin of the ImageJ distribution Fiji. Bar graphs showing percent ciliated cells were plotted in GraphPad Prism and ANOVA was performed to test for statistical significance.

