## Supplemental Figures with Legends for "Mechanistic basis of gene-specific transcription regulation by the Integrator complex"

### SUPPLEMENTAL FIGURE TITLES AND LEGENDS

**Supplemental Figure 1. Titration of 2SC-INTS13ΔC and INTS10-C2S, and mapping of IBMs with corresponding binding sites on INTS13-14.**

(A, B) Western blot of S2C-tagged GFP, INTS13ΔC and INTS10 HEK293-Flp-In cell lines after titration of tetracycline induction. CBP and endogenous proteins were probed to allow judgment of expression levels of the TAP-tagged constructs relative to the endogenous counterpart.

(C, D) Exemplary TAPs from INTS10, INTS13ΔC or GFP cell lines, respectively, that were used for MS analyses. Eluates were visualized on silver-stained SDS-PAGE gels.

(E) Western blot of co-APs of λN-HA-INTS13 or λN-HA-INTS14 with V5-SBP-ZNF608 truncation constructs. V5-SBP-MBP served as negative control.

(F) Coomassie stained SDS-PAGE of co-APs using MBP-ZNF608 constructs that were expressed in *E.coli* and tested for co-purification from lysates with purified 2S-INTS13-14. 2S-GST served as negative control.

(G) Coomassie stained SDS-PAGE of co-APs using GST-INO80E constructs that were expressed in *E.coli* and tested for co-purification from lysates with purified 2S-INTS13-14. 2S-GST served as negative control.

(H) Coomassie stained SDS-PAGE of co-APs using MBP-ZNF655 constructs that were expressed in *E.coli* and tested for co-purification from lysates using purified 2S-INTS13-14. S-MBP served as negative control.

(I- K) Coomassie stained SDS-PAGEs of co-APs using GFP-tagged IBMs that were expressed in *E.coli* and tested for co-purification from lysates with purified 2S-INTS13-14. 2S-GST served as negative control.

**Supplemental Figure 2. Conservation of sIBM sequences of INTS10-13-14 interactors and supplemental information to the INTS13-ZNF655 crystal structure.**

(A) Western blots of co-APs of λN-HA-INTS13 and λN-HA-INO80E as well as endogenous INTS1 with V5-SBP-INO80 from HEK293T cells. V5-SBP was used as negative control.

(B) Sequence alignment of sIBM peptides of ICE2, INO80E, HDGF, ZMYND8, ZNF608, ZNF609, ZEB1, and ZEB2 orthologs from *Homo sapiens* (*Hs*), *Mus musculus* (*Mm*), *Gallus gallus* (*Gg*), *Danio rerio* (*Dr*), *Caenorhabditis elegans* (*Ce*), *Strongylocentrotus purpuratus* (*Sp*) and *Drosophila melanogaster* (*Dm*). Residues with complete conservation (orange) and 70% similarity (light orange) are highlighted. A black box marks the L-x-I-D motif. The β-strand position as observed in the sIBM of ZNF609 is denoted above the alignment. Filled red diamonds indicate amino acids that were mutated in this study. Phosphorylated residues are highlighted in blue.

(C) Domain scheme of TLE3 highlighting a putative sIBM region. Domain are abbreviated as QD – Groucho/TLE N-terminal Q-rich domain, WD40 – WD repeat domain. A sequence alignment of putative sIBM regions for TLE3 orthologs as in (E), and an alignment for human TLE paralogs (TLE1-6) colored as in (B) are also provided.

(D) Western blots of co-APs of endogenous proteins with V5-SBP-TSPYL2 from HEK293T cells. V5-SBP-MBP served as negative control.

(E) Coomassie stained SDS-PAGE of co-APs using MBP-INTS13 domain constructs and GST-ZNF655(IBM) that were co-expressed in *E.coli* and tested for co-purification. GST served as negative control.

(F) Superposition of the INTS13 VWA domain observed in the INTS13 VWA-ZNF655 IBM complex (green and wheat) and in the INTS13-14 structure (grey, PDB ID: 6SN1). The VWA domain of INTS13 is virtually identical in both structures (RMSD: 0.34 Å over 224 aa).

(G) 2Fo-Fc map (blue mesh) of the INTS13 VWA-ZNF655 IBM crystal structure focusing on the ZNF655 hIBM peptide contoured at 0.8σ.

(H) Structure of the INTS13 VWA-ZNF655 IBM complex (green and wheat) superposed onto the INTS13-14 structure (light and dark grey, PDB ID: 6SN1). The ZNF655 contact surface is freely accessible.

(I) Crystal packing of the INTS13 VWA-ZNF655 IBM complex. Surrounding molecules contacting the complex are shown in grey. Since ZNF655 hIBM is found in the same location on all three molecules of the asymmetric unit and is not involved in crystal contacts, its position is unlikely to represent a crystallization artefact.

**Supplemental Figure 3. Supplemental information to the INTS13-14-ZNF609 crystal structure and verification that sIBMs bind the same site on INTS13-14.**

(A) Western blot of co-APs of λN-HA-INTS13 wt or mutant with V5-SBP-INO80E from HEK293T cells. V5-SBP-MBP served as negative control.

(B) Coomassie stained SDS-PAGE of purified 2S-INTS13-14 complex and the 2S-INTS13-14 complex with ZNF609 sIBM fused to the INTS14 C-terminus.

(C) 2Fo-Fc map (blue mesh) of the INTS13-14-ZNF609 sIBM crystal structure focusing on the sIBM peptide (orange) contoured at 0.8σ.

(D) Superposition of INTS13-14-ZNF609 sIBM structure (colored in light and dark green, and orange) with the isolated INTS13-14 structure (grey, PDB ID: 6SN1). Both structures are almost identical (RMSD: 0.30 Å over 1055 residues).

(E, F) Overview (E) and close-up (F) of the surface charge of INTS13-14 from the structure of the complex with ZNF609 sIBM. A positively charged patch on INTS14 α-helical domain that is located directly next to the sIBM binding site is highlighted by a black border.

(F) Same representation as in (E). The ZNF609 peptide is shown as cartoon.

(G) Crystal packing of the INTS13-14-ZNF609 sIBM complex. Surrounding molecules contacting the complex are shown in grey. The INTS13-INT14-sIBM interface is far away from any crystal contacts, making it unlikely to be a crystallization artefact.

(H) Coomassie stained SDS-PAGE of co-APs using MBP-ZNF609 constructs that were expressed in *E.coli* and tested for co-purification from lysates with purified 2S-INTS13-14. 2S-GST served as negative control.

(I) Western blots of co-APs of endogenous INTS13 with wt or mutant V5-SBP-ZEB1 from HEK293T cells. V5-SBP-MBP served as negative control.

(J) Western blots of co-APs of endogenous proteins with wt or mutant V5-SBP-ZMYND8 from HEK293T cells. V5-SBP-MBP served as negative control. Band intensities were quantified and proportional intensities compared to control are listed.

(K, L) Western blots of co-APs of wt or mutant λN-HA-tagged INTS13 or INTS14 with V5-SBP-tagged INTS13 or INTS14 from HEK293T cells. V5-SBP-MBP served as negative control.

(M) Coomassie stained SDS-PAGE of co-APs using MBP-ZNF609 sIBM with 2S-INTS13-14 in the presence or absence of poly-U RNA.

**Supplemental Figure 4. Mapping of IBM from HDGF and verification of INTS13 CMBM binding surface on INTS9-11.**

(A-C) Exemplary TAPs of 2SC-INTS13ΔC or 2SC-GFP from stable HEK293-Flp-In cell lines under different stress conditions that were used for MS analyses. Eluates were visualized on silver-stained SDS-PAGEs.

(D) Coomassie stained SDS-PAGE of co-APs using $His_6$-HDGF constructs that were expressed in *E.coli* and tested for co-purification from lysates with purified 2S-INTS13-14. 2S-GST served as negative control.

(E) Western blots of co-APs of λN-HA-INTS13 wt or mutant with V5-SBP-HDGF from HEK293T cells. V5-SBP-MBP served as negative control.

(F) Affinity measurements by fluorescence polarization of GFP-tagged IBMs and INTS13-14. Error bars indicate standard deviations of means from three experiments.

(G) Published cryo-EM map of INT (EMD-30473-2) at the proposed location of the INTS13 CMBM peptide contoured at 5.5σ with the CMBM model (green) placed inside. Black boxes indicate positions of close-up views in panels (I) and (J).

(H) AF2 model of the INTS9-11 (light and dark blue) complex with INTS13 CMBM (green) (top) and same model colored by pLDDT quality score (bottom).

(I, J) Close-up views of interfaces between INTS9-11 and INTS13 CMBM. Sidechains of interface residues are shown and labeled. Dotted lines represent hydrogen bonds. Residues mutated in this study are underlined. Residues marked by red boxes are mutated in ciliopathy patients.

(K, L) Western blots of co-APs of endogenous proteins or λN-HA-INTS13 with wt or mutant V5-SBP-tagged INTS11 (K) or INTS13 (L) from HEK293T cells. V5-SBP-MBP served as negative control.

**Supplemental Figure 5. Supplemental information for RNAseq and ChIPseq experiments.**

(A, B) Western blot validations of siPOOL knockdowns of INTS13, HDGF, TSPYL2, ZEB1, ZMYND8, ZNF609 and ZNF687, and RT-qPCR validation of siPOOL knockdowns of ZNF655 and ZNF608 in HEK293T cells used for RNAseq replicates 1 (A) and 2 (B).

(C) GO-term enrichment analysis of DEGs common in INTS13 and individual TF knockdowns. Significantly enriched terms ($p < 0.05$) are ordered by decreasing number of associated genes.

(D) Heatmaps of INTS13 (left) and INTS11 (right) ChIP signals at INTS13 peaks centered around TSSs ±5 kb.

(E) Venn diagram of the overlap between INTS13 and INTS11 bound genes in ChIP experiments.

(F) Pie charts representing the proportions of INTS13 and INTS11 promoter-bound genes in INTS13 and INTS11 ChIP, respectively.

(G) Heatmaps of INTS13 (left) and INTS11 (right) ChIP signals at INTS13 bound genes ±5 kb around TSSs.

(H) Density profiles of INTS13 (top) and INTS11 (bottom) ChIP signals at INTS13 promoter-bound (green) and non-promoter bound (olive) genes.

(I) Venn diagram of the overlap between INTS13 and INTS11 promoter-bound genes.

(J) Heatmaps of INTS11 ChIP signals ±1 kb around HDGF (left) or ZEB1 (right) peak summits.

(K) Venn diagrams of overlap between INTS13 promoter bound genes (green) and genes bound by HDGF (orange), ZEB1 (red) and ZNF609 (yellow).

(L) Venn diagram of INTS13 promoter bound genes (light green) and INTS13 DEGs (dark green).

(M) Venn diagram of genes not bound by INTS13 at their promoters (olive) and INTS13 DEGs (dark green).

**Supplemental Figure 6. Supplemental data to RNAseq after glucose starvation and to IFs.**

(A-C) Western blot validation of siPOOL depletions of INTS13 (A), HDGF (B) and ZMYND8 (C) under starvation conditions in HEK293T cells used for RNAseq replicates.

(D, E) Volcano plots of RNAseq experiments showing L2FCs and $p_{adj}$ of transcript levels under ctrl siPOOL (D) or INTS13 siPOOL (E) treatment upon glucose starvation relative to normal growth conditions (n=2 per condition). Significant DEGs (|FC| > 1.5, $p_{adj} < 0.05$) are colored red for up- and blue for downregulated transcripts. Number of transcripts in each category is noted above the graph.

(F-I) Scatter plots showing $\log_{10}$ of normalized read counts of transcripts under glucose starvation versus normal growth conditions with ctrl siPOOL treatment. Different classes of genes are highlighted. (F) DEGs upon glucose starvation (fuchsia), (G) INTS13 dependent (green) and INTS13 independent (purple) starvation response genes, (H) INTS13 dependent starvation response genes that were upregulated under normal growth conditions upon INTS13 siPOOL depletion (moss green), and (I) INTS13 dependent starvation response genes that were downregulated under normal growth conditions upon INTS13 siPOOL depletion (light green).

(J, K) Venn diagrams of the overlap between DEGs upon glucose starvation in ctrl and HDGF (J) or ctrl and ZMYND8 (K) siPOOL treated cells.

(L) Venn diagram of the overlap between INTS13-dependent starvation response genes that were upregulated upon INTS13 depletion (moss green) and all HDGF (orange) and ZMYND8 (red) dependent starvation response genes.

(M) Western blot validation of siPOOL depletions of INTS13, ZMYND8, HDGF. ZNF609 and ZNF687 in HEK293T cells corresponding to Figure 6 (H-J).

(N, O) Western blot validation of siRNA depletion of endogenous INTS13 (N) and HDGF (O) and expression of siRNA resistant V5-SBP-INTS13 or HDGF rescue constructs in HEK293T cells corresponding to Figure 6 (K-N).

**Supplemental Figure 7. Biochemical validation of completed INT model.**

(A) Coomassie stained SDS-PAGE of co-APs of INT subcomplexes with purified GST-INTS15 from lysates of insect cells expressing human INT subunits. 2S-GST served as negative control.

(B) Coomassie stained SDS-PAGE co-APs of a minimal co-expressed INTS5-10-15 complex in insect cells using affinity tags on MBP-INTS10 (1-54) (Amylose), 2S-INTS5 (1-234) (StrepTactin) or GST-INTS15 (Glutathione).

(C) AF2 model of the minimal INTS5-10-15 complex (red, purple, and pink). Black boxes mark positions of close-up views of interfaces shown in panels (E) and (F).

(D) AF2 model of the INTS5-10-INTS15 complex from panel (C) colored by pLDDT quality score.

(E, F) Close-up views of the modelled INTS5-15 and INTS10-15 interfaces. Sidechains of interface residues are shown and labeled. Residues mutated in this study are underlined.

(G, H) Western blots of co-APs of endogenous proteins with wt or mutant V5-SBP-tagged INTS5 or INTS10 from HEK293T cells. V5-SBP-MBP served as negative control.

(I) AF2 model of the complex between INTS10 (pink) and INTS14 VWA (light green). The black box marks the position of the close-up view of the complex interface shown in panel (K).

(J) AF2 model of the INTS10-14 VWA structure from panel (I) colored by pLDDT quality score.

(K) Close-up view of the INTS10-14 VWA complex interface. Sidechains of interface residues are shown and labeled. The putative position of the $Mg^{2+}$-ion is indicated. Residues mutated in this and our previous study[1] are underlined.

(L) Superposition of the VWA domain of ITGAL (light grey) and its interacting protein ICAM3 (dark grey) (PDB ID: 1T0P)[2] on the INTS10-14 VWA complex model highlights similarities of ligand coordination in the MIDAS pocket.

(M) CryoEM map of the published INT structure (EMD-30473-3) around the proposed position of DSS1 contoured at 9.0σ with DSS1 model inserted. INTS2 (dark grey), and INTS7 (light grey) contact DSS1 (yellow).

#### Supplemental Figure 1

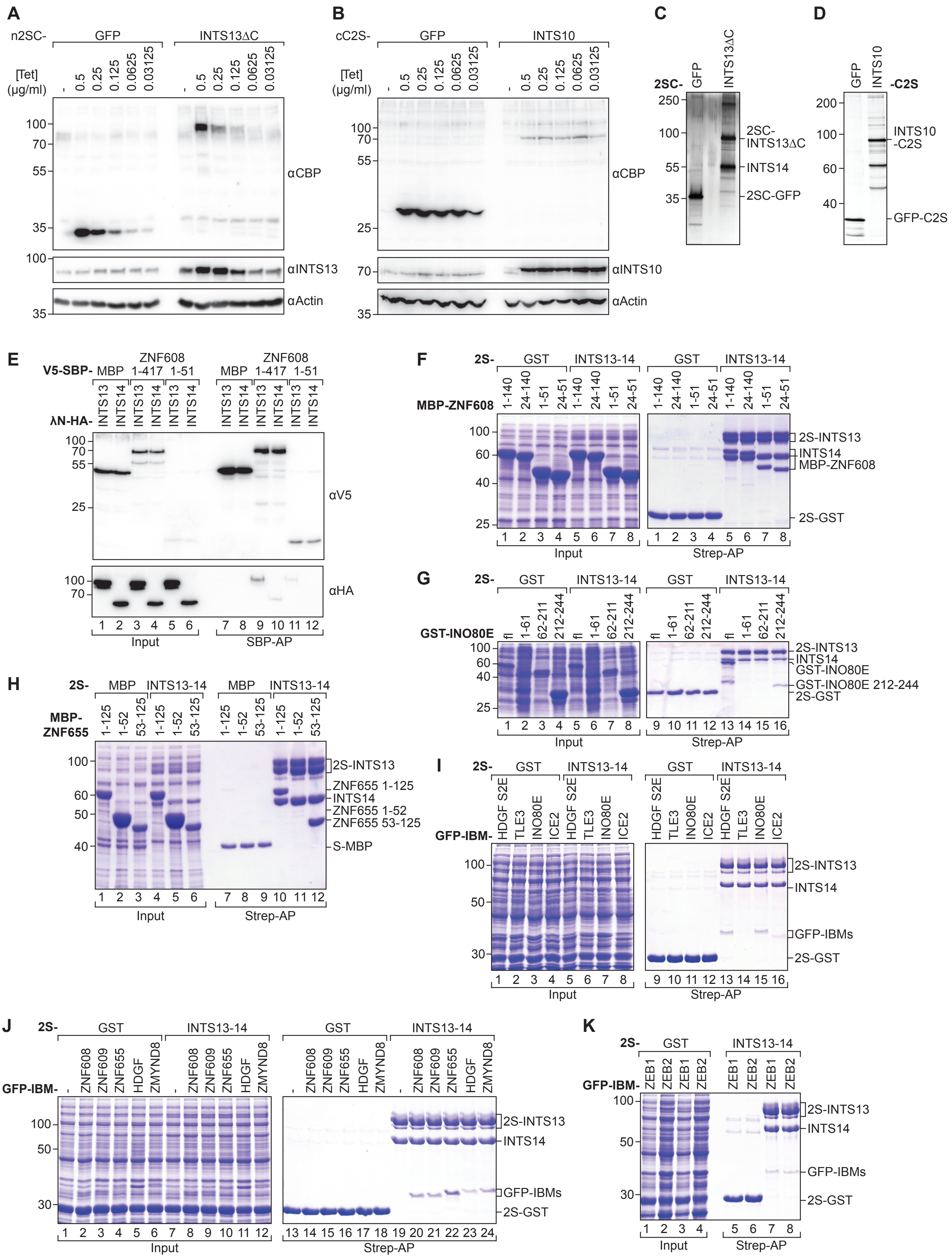

#### Supplemental Figure 2

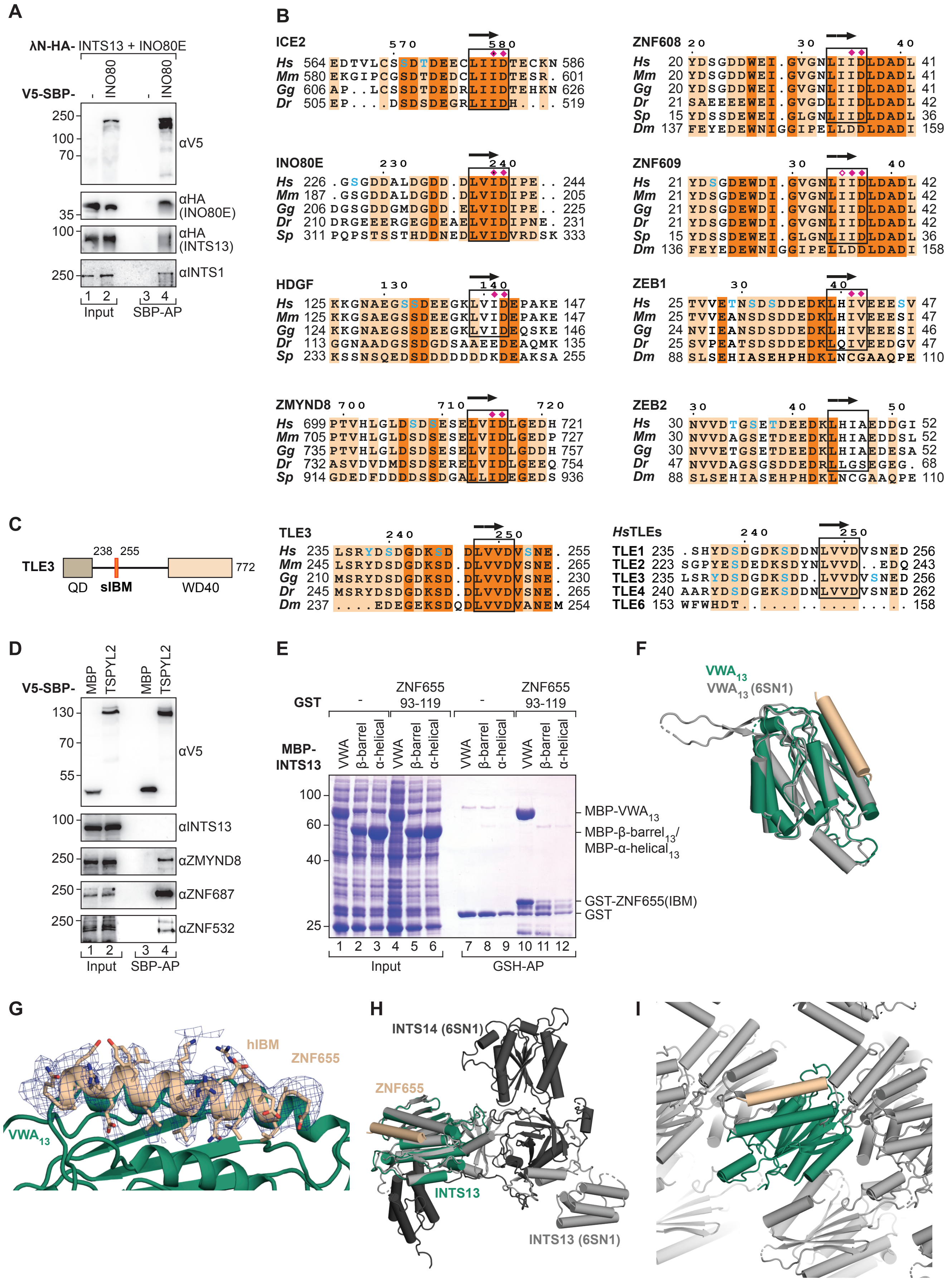

### Supplemental Figure 3

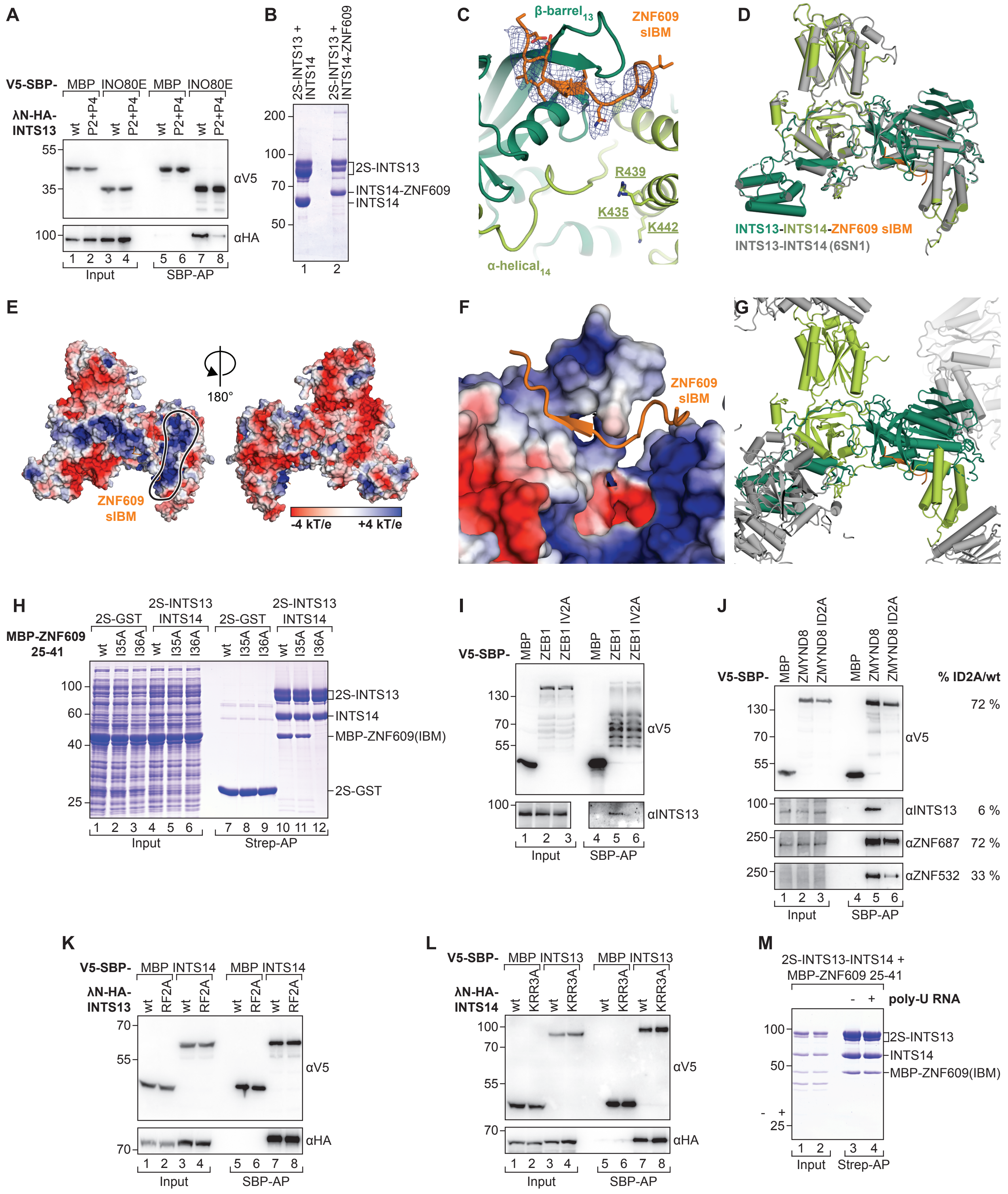

#### Supplemental Figure 4

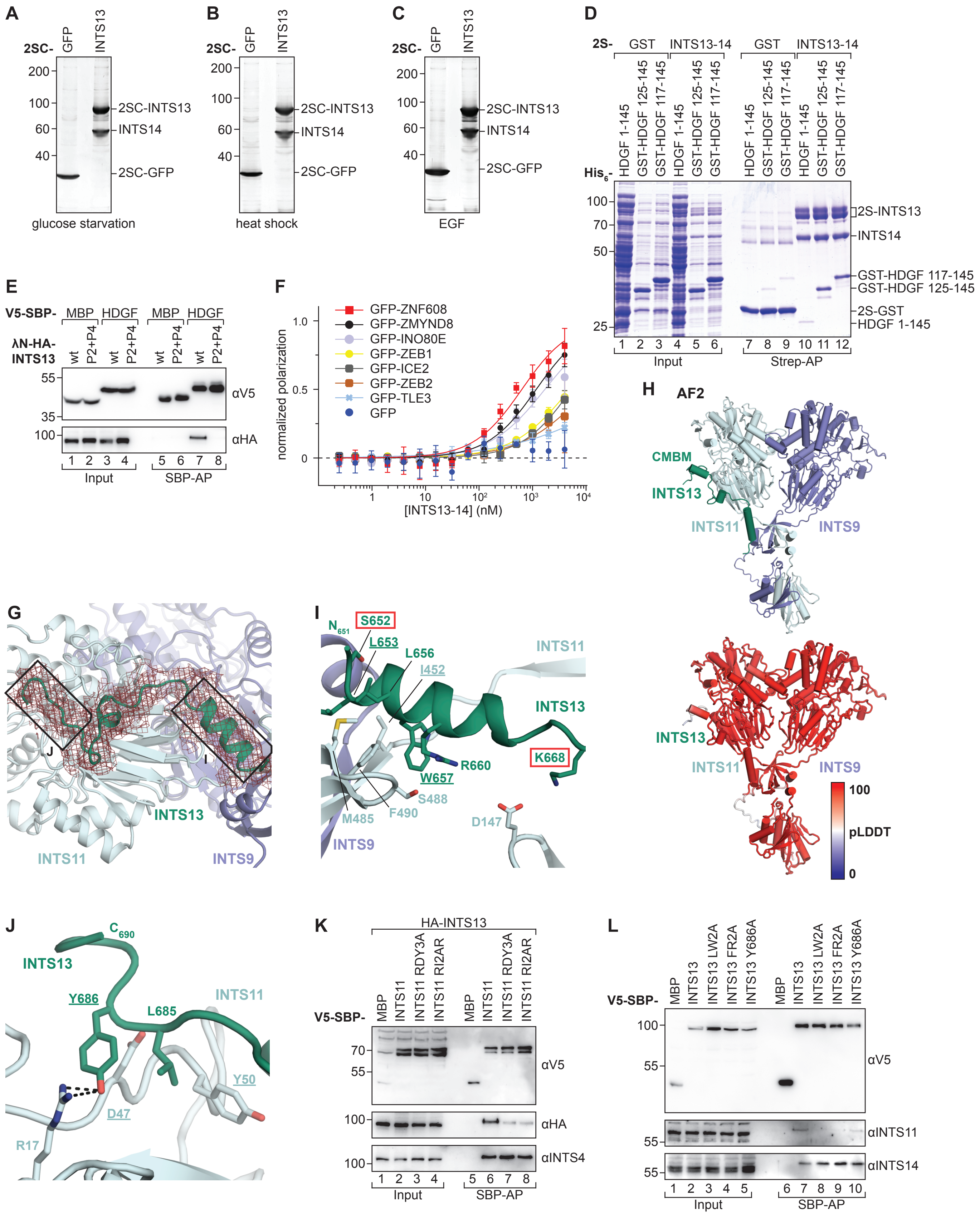

### Supplemental Figure 5

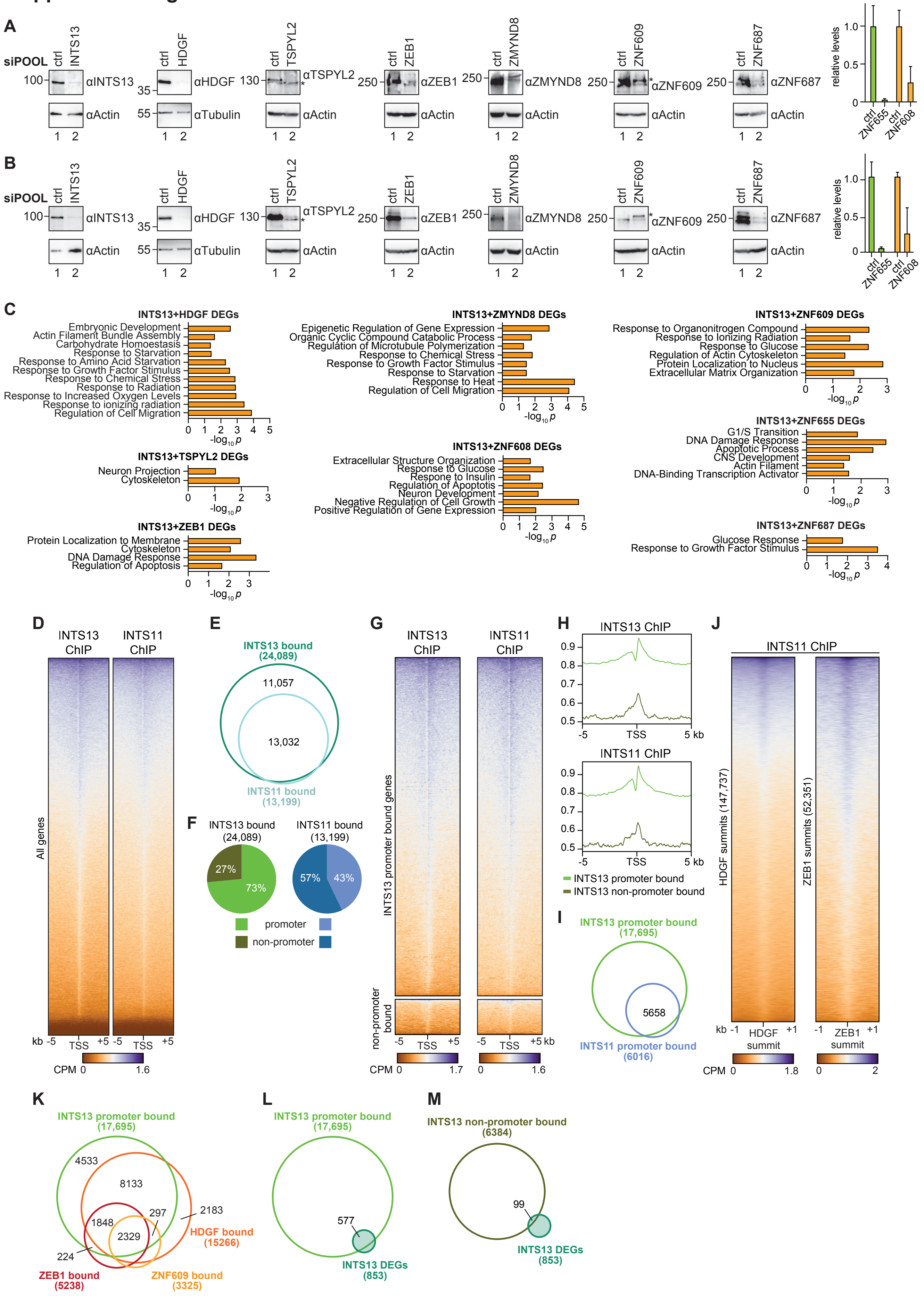

#### Supplemental Figure 6

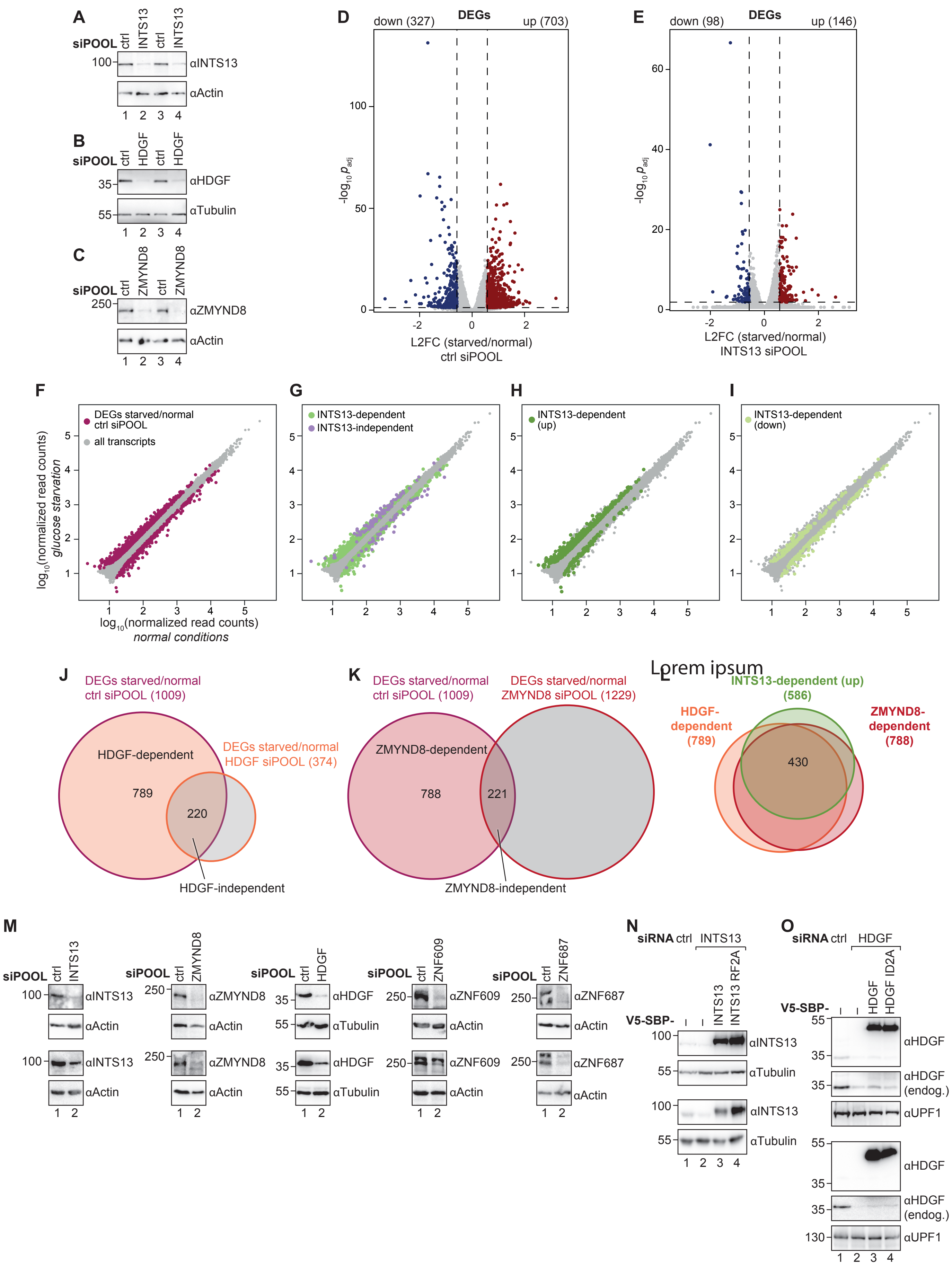

#### Supplemental Figure 7

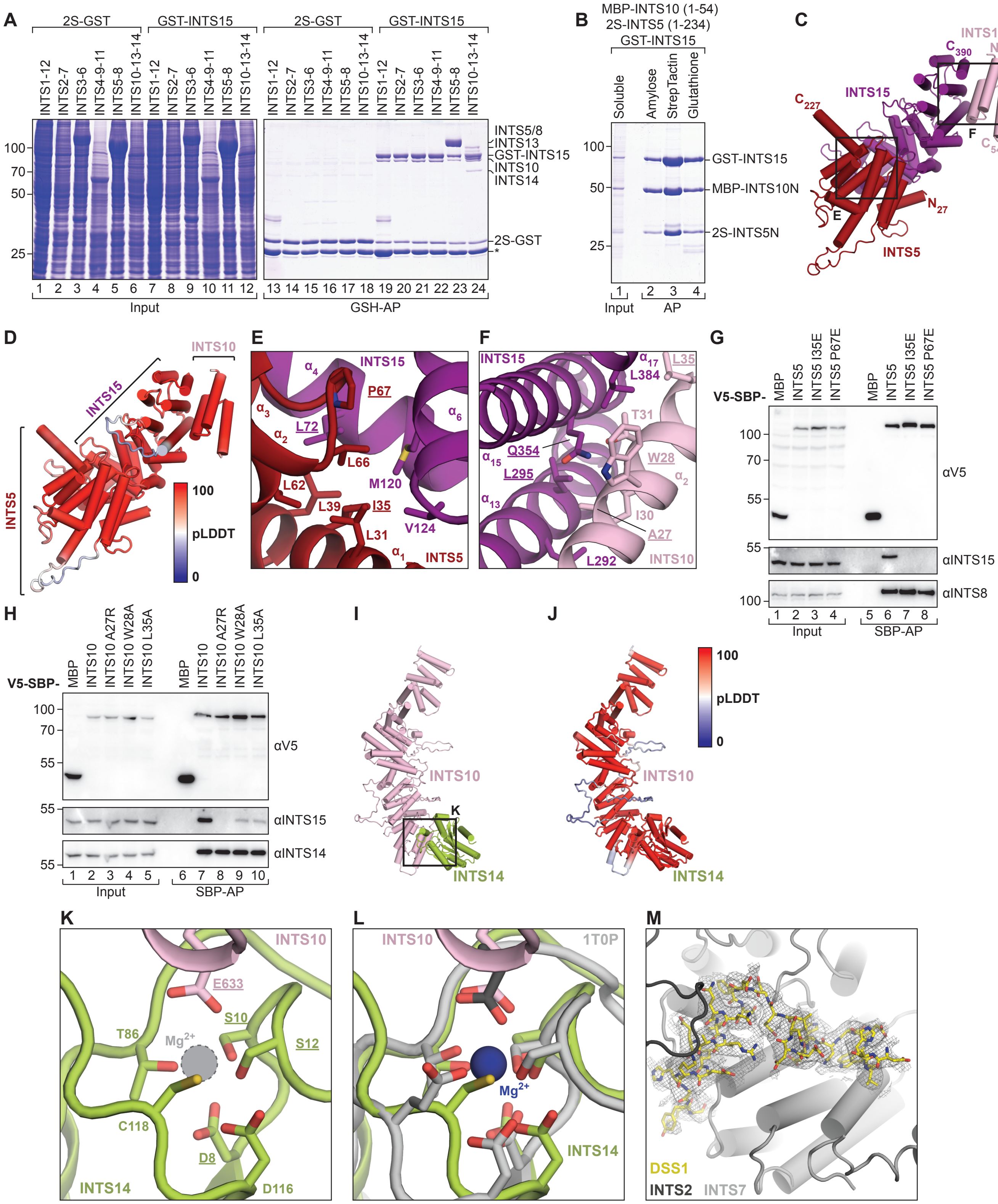
